# Spatial determinants of tumor cell dedifferentiation and plasticity in primary cutaneous melanoma

**DOI:** 10.1101/2025.06.21.660851

**Authors:** Tuulia Vallius, Yingxiao Shi, Edward Novikov, Shishir M Pant, Roxanne Pelletier, Yu-An Chen, Juliann B. Tefft, Ajit Nirmal Johnson, Zoltan Maliga, Guihong Wan, Mia DeSimone, George Murphy, Sandro Santagata, Yevgeniy R Semenov, David Liu, Christine G Lian, Peter K Sorger

## Abstract

Localized cutaneous melanoma can be cured by excision but success critically depends on early detection and risk assessment of primary lesions. However, their initiation, progression, and immunology remain poorly understood, partly due to high intra- and inter-tumor heterogeneity. We studied this heterogeneity using spatial profiling in over 300 histological domains, each representing a single progression stage, and found that 200-600 cell neighborhoods from a single melanoma can be as different in RNA and protein expression as neighborhoods from different tumors. These differences are not stochastic, however, and disease progression can be mapped at the neighborhood level onto a cell state landscape defined by the activity of the melanocyte master regulator MITF and genes associated with a dedifferentiated neural crest phenotype. Position in this landscape is influenced by proximity to immune cells, perivascular environments, and other tissue features, but no single association is absolute, giving rise to complex spatial patterns.

## INTRODUCTION

Cutaneous melanoma is the most aggressive type of skin cancer: metastatic disease remains lethal in many patients, despite the approval of targeted therapies and immune checkpoint inhibitors (ICIs).^1^ The incidence of cutaneous melanoma is projected to double between 2020 and 2040 due to changes in sun exposure and diagnostic criteria^2^ but local surgery can be curative if the disease is detected early. Early detection is dependent on routine skin surveillance^3^ followed by biopsy or excision of suspicious lesions for histopathological evaluation. This involves examination of hematoxylin and eosin (H&E) stained tissue sections prepared from formalin-fixed paraffin embedded (FFPE) specimens, complemented in some cases by immunohistochemistry (IHC) but rarely by sequencing. Decades of H&E and IHC-based histopathology have led to highly refined morphological criteria for melanoma diagnosis and staging^4^ but identifying aggressive tumors remains difficult and a substantial portion of metastatic disease involves recurrence in patients who have undergone surgery with curative intent.^5^ This inability to predict melanoma recurrence and the countervailing pressure to avoid overdiagnosis^6^ highlights a substantial need for new approaches to predicting melanoma risk based on better understanding of cellular and molecular properties associated with progression in primary disease. Fortunately, the ready availability of FFPE specimens representing the earliest stages of melanoma and the introduction of spatial profiling methods well-suited to the limitations of these small specimens represent an excellent setting in which to study how solid tumors first arise and overcome immune surveillance.

The initiation and progression of cutaneous melanoma is a multistep molecular process^7–11^ involving distinct histological stages: (i) a premalignant state (either atypical melanocytes in “precursor fields” or nevi), (ii) *melanoma in situ* (MIS), a pre-invasive state, (iii) locally invasive cancer typically exhibiting a radial growth phase followed by a deeply invasive vertical growth phase (RGP and VGP), (iv) locoregional metastasis (e.g., to draining lymph nodes), and (v) distant metastasis to the skin, brain, lungs, and liver. In some specimens, multiple stages of primary melanoma from precursor field to invasive tumors can be found in a single tissue block, creating an opportunity to study progression in a single individual.^12^

DNA sequencing has established an essential role for driver mutations in oncogenes such as *BRAF*, *NRAS* and *KIT* during melanoma initiation and metastasis,^13–16^ and single-cell sequencing and IHC have demonstrated a role for lineage plasticity and tumor cell dedifferentiation in therapy resistance, particularly in the metastatic setting.^17–24^ MITF is a master regulator and critical factor in melanocyte differentiation, melanosome formation, and melanin biosynthesis and genes controlled by *MITF* can be either up- or down-regulated during melanoma progression. For example, during induction of “epithelial-to-mesenchymal” transitions (EMT) in progressing melanomas, *MITF* expression decreases,^25^ leading to repression of transcriptional programs involved in pigment production. Inflammatory microenvironments have been reported to induce cell plasticity^26–29^ and a switch from *MITF*-driven melanocytic states to an invasive dedifferentiated state characterized by expression of *NGFR*, *AXL*, and *SOX9*^13,18–20,30^. The presence of endothelial cells is another factor proposed to contribute to dedifferentiation and melanoma growth (in mouse models).^31^

Both primary and metastatic melanoma are characterized by high intra-tumoral heterogeneity^32,33^ and this has direct implications for the use of biomarkers in diagnosis.^34–36^ Heterogeneity within a single specimen^37^ and across specimens of similar stages^36^ in the expression of *MITF*-regulated proteins such as MART1^38^ and PMEL^39^ or the SOX10 transcription factor^40^ has made it challenging to identify a definitive progression biomarker or biomarker combination. Thus, no molecular or genetic features are routinely part of diagnosis in the primary disease setting. Instead, the current melanoma staging system (the AJCC 8th edition for melanoma staging)^41^ relies primarily on visual assessment of H&E-stained sections to score the depth of invasion (the Breslow depth), presence or absence of ulceration (which can also have a macroscopic presentation), and whether or not lymph node metastases are present.^4,41^ The presence of tumor-adjacent lymphocytes (a “brisk” TIL response)^42^ is often recorded in pathology reports but the prognostic significance remains uncertain.^43^

A further limitation in our understanding of primary melanomas is that molecular profiling^44^ has been performed primarily in metastatic disease^13,14,33^, with primary disease remaining less thoroughly studied.^12^ In part, this is because formaldehyde fixation and small specimen size (along with a requirement to preserve tissue for patients who are still alive) make traditional RNA and scRNA sequencing difficult.

Studies (including those from our own laboratories) on limited numbers of primary melanomas have shown that EMT-like transcriptomic programs are associated with poor survival in primary melanoma^25^, and that de-differentiation can be observed in pre-treatment and pre-metastatic stages.^12,25,45–47^ These studies suggest that additional genetic and epigenetic changes found in metastatic disease might also be present in primary melanoma.

In this paper we characterize inter- and intra-tumoral heterogeneity of primary melanomas at a molecular level with the goal of distinguishing random variation (as observed for passenger mutations in metastatic cancers)^48^ from recurrent and patterned variation in over 300 distinct histological domains from 62 primary cutaneous melanomas. The majority of these specimens contained multiple progression stages ranging from adjacent normal skin to the cutaneous component of Stage III (metastatic) melanoma.

Patterns of protein and gene expression were measured using cyclic multiplexed immunofluorescence (CyCIF)^49^ imaging and GeoMx microregional transcriptomics.^50^ Confirming previous histological and IHC data, we failed to identify any single protein feature predictive of disease stage and found that microregions (MRs; typically 200-600 histologically similar tumor cells) from a single specimen could differ from each other to as great a degree as MRs from all specimens. However, these differences were not random but were instead organized spatially based on MITF activity, melanocyte dedifferentiation, and other molecular features. Moreover, these features were strongly associated with the degree of local inflammation and proximity to the vasculature. Thus, both inter- and intra-tumoral heterogeneity in primary melanoma appear to be influenced in large part by the local microenvironment, which is in turn patterned by mesoscale features such as vessels and lamina that organize normal tissue.

## RESULTS

### Patient cohort and spatial data generation

Archival FFPE specimens of primary cutaneous melanoma from 62 patients (**Fig. 1A,B**; MEL14 to MEL86; see **Supplementary Table S1** for HTAN identifiers) were reviewed by a board-certified dermatopathologist and annotated for the presence of five histological features associated with disease progression: (i) adjacent normal skin, (ii) melanocytic atypia (precursor fields), (iii) melanoma in situ (MIS), (iv) radial growth phase (RGP) and (v) vertical growth phase melanoma (VGP; **Fig. 1C,D; Supplementary Fig. S1A**). Sample set A (n=45) was selected for the presence of multiple progression stages within the same sample with an emphasis on stage I and set B (n=17) was selected for stage II disease; multiple stages were often present in these specimens, but this was not a selection criterion.

**Figure 1.**
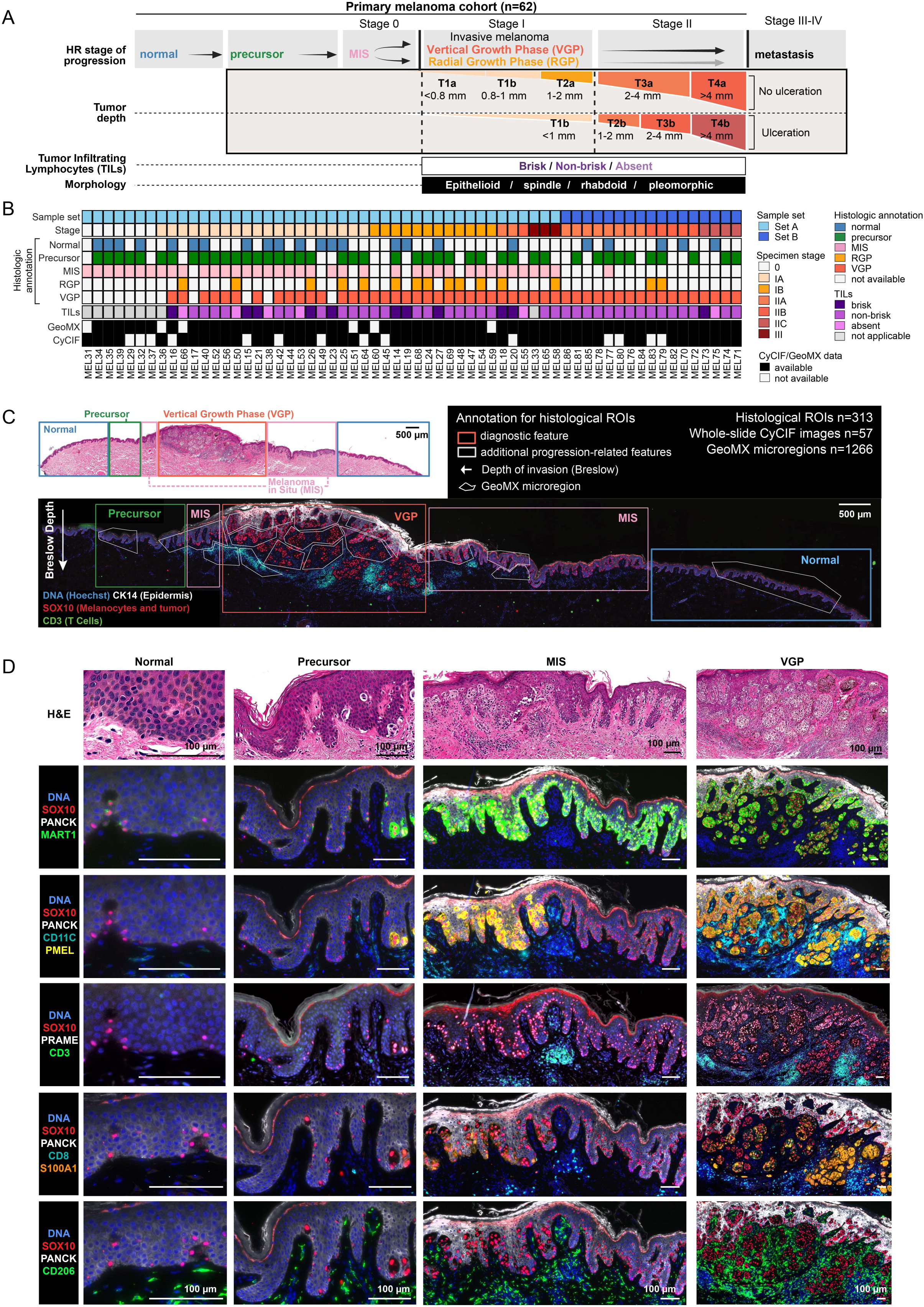
Multimodal profiling of primary melanoma. **A.** Summary of the current AJCC 8th edition for cutaneous melanoma staging^41^ based on features identified from H&Es. **B.** Overview of the primary melanoma cohort used for multimodal spatial profiling and their associated clinical (AJCC) and histological characteristics. (TILs: Tumor-infiltrating lymphocytes; normal skin; precursor; MIS: melanoma in situ; RGP: radial growth phase melanoma; VGP: vertical growth phase melanoma). **C.** Example specimen (MEL14) annotated with clinical histological stages (colored rectangles), as H&E (top) and CyCIF image of a serial section (bottom). CyCIF image shows a subset of markers: DNA (blue), CK14 (epidermis; white), SOX10 (melanocytes and tumor cells; red), and CD3 (T cells; green). White polygons overlaid represent the microregions (MRs) used for spatial transcriptomics (GeoMx). Arrow indicates the measurement and direction of Breslow depth. Scale bars, 500 μm. **D.** Examples of histopathological features annotated in the specimen MEL14. Top row: Fields of view of a H&E-stained section with four major histologic regions indicated: adjacent normal melanocytes and skin, melanocytic atypia (precursor), melanoma in situ (MIS), vertical growth phase melanoma. Bottom rows: Corresponding CyCIF images showing staining for a variety of tumor and immune markers. See **Supplementary Table S3** for how markers related to cell types. Scale bars, 100 μm.

Melanocytic nevi and *de novo* melanomagenesis from precursor fields (morphologically atypical melanocytes^51^) are the two currently recognized routes to melanoma initiation^7^ with current evidence suggesting that nevi initiate ∼20% of melanomas and precursor fields ∼80%.^52,53^ This study focuses on the latter: precursor fields comprising discontinuous, non-pagetoid intraepidermal proliferation of cytologically dysplastic melanocytes located at the perimeter of, and in contiguity with, stage 0-III melanoma. Upon progression, these atypical melanocytes continuously replace the basal layer of the epidermis and/or exhibit pagetoid scatter giving rise to MIS (Stage 0). Stages I & II melanoma involve increasing depths of dermal invasion (Breslow depth) and Stage III includes locoregional metastasis (**Supplementary Fig. S1A**).^54,55^ Histopathological annotation of 62 specimens yielded a total of 313 distinct histological regions (HRs) each representing a different progression stage (ranging from 1 to 14 per specimen; average 4); these HRs were also scored for the presence of tumor infiltrating lymphocytes (TILs), and for epithelioid, spindle, rhabdoid, and pleomorphic tumor cell morphologies.

We then performed 34 to 49-plex whole-slide CyCIF imaging on these specimens (**Supplementary Fig. S1B**) and used histopathological annotations and CyCIF data to guide the selection of microregions (MRs) for GeoMx microregional transcriptomic profiling. These MRs ranged in size from ∼100 cells for normal and precursor areas to ∼350 cells for RGP and VGP regions (**Fig. 1C; Supplementary Fig. S1C,D**) which is similar to the correlation length scales for many tumor features, as previously described.^56^ Prior to mRNA preparation, MRs containing a mixture of melanocyte/tumor cells and other cell types were separated into tumor, non-tumor and immune compartments using either a mixture of anti-SOX10 and anti-MART1 antibodies or anti-CD45 antibodies (referred to as “segmentation” in the GeoMx workflow; **Supplementary Figs. S1D, S2A** and **Supplementary Table S2**); a total of 1266 MRs were profiled. Archival specimens of primary melanoma can be difficult to manage when friable; thus, not all specimens that generated GeoMx and CyCIF data passed quality control and could be included in the final dataset (**Fig. 1B**).^57^

To identify protein features associated with melanoma initiation and progression, CyCIF images were segmented using algorithms in MCMICRO^58^ and marker intensities were quantified in ∼2 x 10^6^ tumor and ∼3 x 10^6^ non-tumor cells followed by gating and cell type calling (see **Supplementary Fig. S2B-E** for markers and cell type identification plan). Melanocytes and tumor cells were further subdivided based on the expression of melanocytic lineage markers (e.g., MART1, S100A1), clinical biomarkers (e.g., PRAME), differentiation markers (e.g., SOX9, SOX10, NGFR), and proliferation markers (e.g., KI67, pH3; see **Supplementary Fig. S2C** for gating strategy and **Supplementary Table S3** for index of gene and protein names). The proportions of tumor cells corresponding to melanocytic, transitional, mesenchymal, and neural-crest (NC) lineages were then determined based on patterns of SOX10, MART1, SOX9, and NGFR marker co-expression (**Fig. 2A,B**).

**Figure 2.**
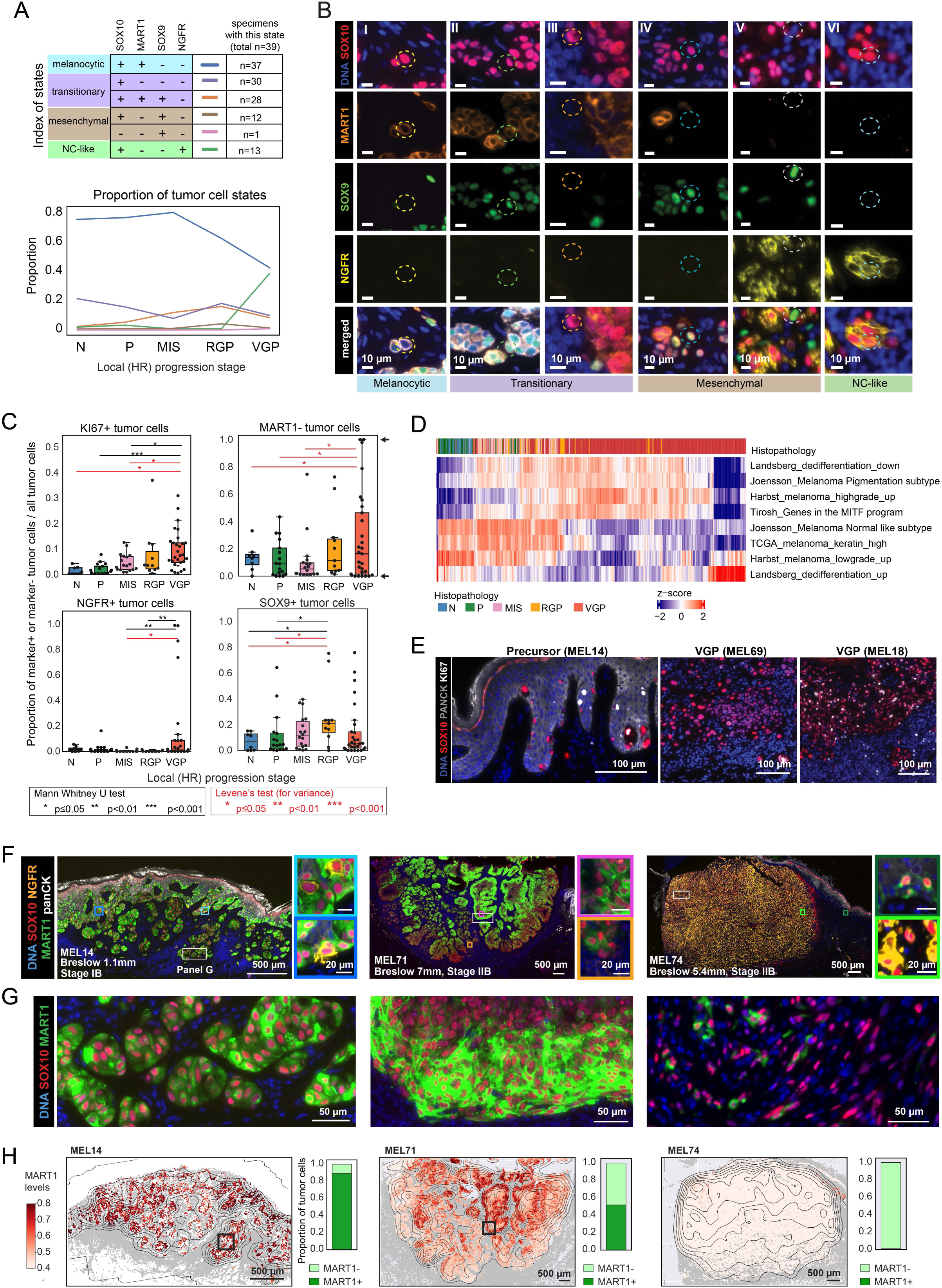
Association of melanocytic differentiation with progression and intertumoral heterogeneity. **A.** Tumor cell states annotated according to the binarized staining of protein markers for SOX10, MART1, SOX9 and NGFR (upper panel). The mean proportion of each melanocyte and tumor cell state within local stages of progression (N, P, MIS, RGP and VGP; lower panel). **B**. CyCIF images showing examples of single cells representing the protein-based tumor cell states detected in the primary melanoma cohort samples. The magnified regions labeled as I-VI from **Supplementary Fig. S3B**, stained for DNA (blue), SOX10 (red), MART1 (orange), SOX9 (green) and NGFR (yellow). Scale bars, 10 μm. **C.** The proportion of SOX10+ melanocytes or tumor cells expressing KI67, MART1, NGFR and SOX9 grouped by histologic regions (HR). Each data point represents a sample proportion, calculated from a single sample. MannWhitney U test (black) and Levene’s test (red) *, *P* < 0.05; **, *P* < 0.01; ***, *P* < 0.001. Box represents the first and third interquartiles of the data, whiskers extend to show the rest of the distribution except for points that are determined to be outliers. See **Supplementary Table S5** for the CyCIF quantification. **D.** Heatmap showing mean expression (z-scored) of melanoma associated signatures within melanocyte and tumor GeoMx MRs annotated for local stages of progression. **E**. CyCIF images of specimen MEL14, MEL69 and MEL18, showing selected fields of view stained for DNA (blue), SOX10 (red), KI67 (white), PanCK (gray). **F**. CyCIF images of specimens MEL14 (left), MEL71 (middle) and MEL74 (right), showing a subset of markers: DNA (blue), epidermis (panCK: white), and melanocyte/tumor (SOX10: red, NGFR: yellow, MART1: green). Colored rectangles represent magnified regions at right. Scale bars, 500 μm and 20 μm. **G.** Selected fields of view of CyCIF images from panel F (white rectangles), showing DNA (blue), SOX10 (red), and MART1 (green). Scale bars, 50 μm. **H.** Scatter plots showing mean MART1 fluorescence intensity for tumor cell and melanocyte cell centroids for invasive regions from MEL14, MEL71 and MEL74. The centroids for other cell types (non-tumor cells) are shown with grey dots. The proportion of MART1^+^ and MART1^-^ tumor cells (binarized call with gate 0.5) is shown on the right of each scatter plot. Scale bars, 200 μm.

### Association of melanocytic differentiation with progression and intertumoral heterogeneity

The majority of melanocytes in normal and precursor regions were melanocytic (defined as SOX10^+^ MART1^+^, **Fig. 2A-C**, **Supplementary Fig. S3A,B, Supplementary Table S4**). Mesenchymal states (MART1^-^ and SOX9^+^) emerged only in invasive melanomas, and de-differentiated neural-crest like tumor cells (NGFR^+^) were restricted to later progression stages (**Fig. 2A,C**). Similar trends were observed for Breslow depth, a measure of the maximal depth of dermal invasion used for clinical staging (**Supplementary Fig. S3C**). These results are consistent with an extensive histopathology literature based on H&E staining and single channel IHC of primary melanoma.^34–36^ When GeoMx transcriptomic profiles from 900 tumor MRs were compared to published transcriptional signatures derived from scRNA sequencing of metastatic tumors and melanoma cell lines^26,59,60^, a good correspondence was observed to local progression stage (**Fig. 2D**). For example, the “*Harbst melanoma highgrade*” and “*Joensson_Melanoma Pigmentation subtype*” signatures were enriched in RGP and VGP relative to normal, precursor and MIS stages, whereas the opposite was true for “*Harbst melanoma lowgrade*” and “*Joensson_Melanoma Normal like subtype*”. Thus, local progression in primary melanoma specimens was associated with the emergence of dedifferentiated states that are similar to states previously characterized in metastatic disease.

Despite the correspondence between the average disease stage and the average protein or gene expression, CyCIF revealed a high level of specimen-to-specimen difference in the proportions of cells positive for tumor state markers such as MART1, NGFR, SOX9, PRAME and S100A1 or KI67^+^ at each stage (**Fig. 2C**, **Supplementary Fig. S3D,E**). For example, the average proportion of proliferative tumor cells (a feature predictive of poor prognosis^61,62^) increased significantly with stage, but a subset of invasive VGP HRs were as nonproliferative in the tumor compartment (2% Ki67^+^) as the average precursor field (**Fig. 2C,E**). Similarly, the proportion of MART1^-^ cells in VGP HRs varied from 0% to 100% (**Fig. 2C,H**). MART1 (*MLANA*) levels are controlled by the melanocyte master transcription factor MITF^63^ and loss of MART1 expression reflects a fundamental change in tumor differentiation status (**Supplementary Fig. S3F**); MART1 staining is also widely used in melanoma diagnosis.^64^ However, we found that it was not the mean proportion of MART1^-^ cells that increased significantly with progression stage but rather the degree of intertumoral variability (as measured by Levene’s test; **Fig. 2C**). The proportion of NGFR^+^ tumor cells exhibited a similar trend of increasing variance with stage (e.g., MIS to VGP stages; Levene’s test p=0.05, **Fig. 2C**). Note that this type of analysis underestimates the extent of cell-to-cell difference since it binarizes marker levels into plus and minus states and ignores evidence of continuous gradation in expression both within whole specimens and within individual domains of a single specimen (e.g. of MART1; **Fig. 2F-H, Supplementary Fig. S3G, S5H**).

### Gene expression programs associated with intra and inter-tumor heterogeneity

To investigate the origins of inter- and intra-tumor heterogeneity, we analyzed gene expression in 900 GeoMx tumor and melanocyte MRs using Principal Component Analysis (PCA). This revealed a remarkably well organized trajectory in PCA space from normal HRs (**Fig. 3A** light blue data points) to invasive RGP and VGP HRs on the right (red data points) with extensive intermixing of patient IDs (**Fig 3B-C; Supplementary Fig. S4A-E**)^13^. When the PCA landscape was annotated using previously established gene expression signatures (e.g. from **Fig. 2D**), we found that MRs lying to the left of PC1 corresponded to normal-like melanoma signatures (“*Joensson_Melanoma Normal”)* and those on the right to dedifferentiated signatures (“*Landsberg_dedifferentiation_up*”, **Fig. 3D,E; Supplementary Fig. S4F; Supplementary Fig. S5A-D, Supplementary Fig. S6A-C**). During embryogenesis, melanoblasts arise from the neural-crest lineage and migrate to the epidermis where they mature into melanocytes.^65^ It has been proposed that melanocytes expressing neural crest markers, NGFR in particular, represent melanoma initiating cells or cancer stem cells.^66^ In our data embryogenic signatures (*Gopalan_Mesenchymal_Early*)^67^ were strongly enriched to the right of PC1, coincident with neural crest signatures (e.g. “*Rambow_Neural Crest Stem Cell*”) and TGF-β pathway activation (**Supplementary Fig. S4G-J**), all of which have previously been associated with dedifferentiated melanocyte states.^68,69^ Separation along PC2 corresponded to the strength of the MITF signature: MITF-intermediate states were observed in normal HRs (green outline, **Fig. 3F,G**, **Supplementary Fig. S4G**), MITF-high (“hyper-pigmented”) states in invasive melanoma (red outline), and MITF-low dedifferentiated neural-crest like states in invasive VGP HRs (blue outline). Separation of MRs along PC2 is consistent with MITF functioning as a molecular rheostat: MITF-intermediate states were observed in normal HRs (green outline, **Fig. 3F,G**, **Supplementary Fig. S4G**), MITF-high (“hyper-pigmented”) states in invasive melanoma (red outline), and MITF-low dedifferentiated neural-crest like states in invasive VGP HRs (blue outline). Thus, MITF activity and tumor cell dedifferentiation emerged as two key features organizing tumor states. In contrast, neither the solar elastosis score (as evaluated at the level of whole specimens) nor the strength of UV transcriptional signatures (at the level of individual MRs) exhibited systematic variation along PC1 or PC2; thus, UV exposure did not appear to be a primary determinant of local tumor cell state (**Fig. 3I**, **Supplementary Fig. S7A-D**) and this was also true of the interferon-γ (IFNγ) or IFNα response signatures (**Supplementary Fig. S4K**, **L**).

**Figure 3.**
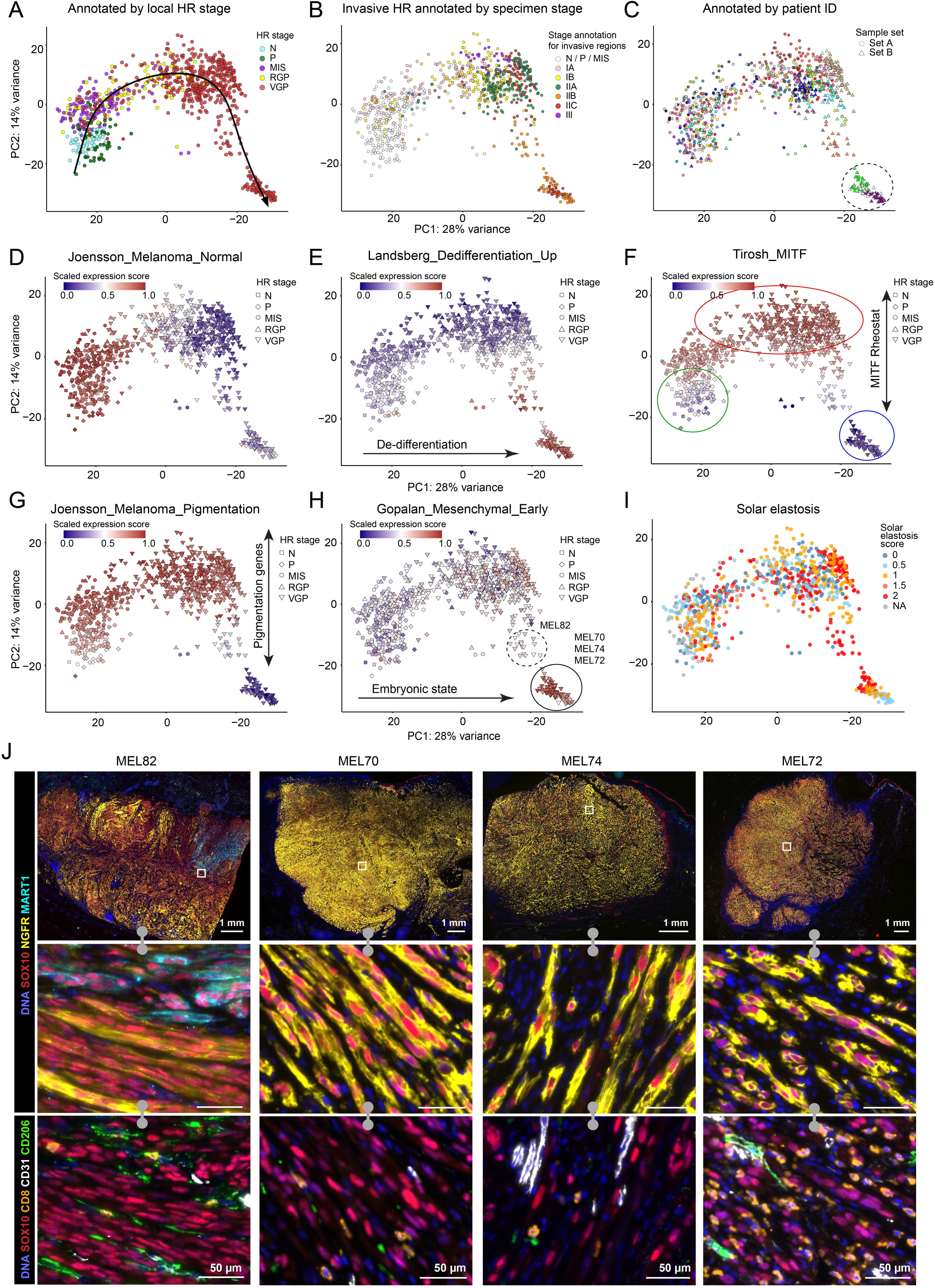
Intra- and inter-tumoral transcriptomic heterogeneity in primary melanoma. **A-C.** Principal component (PC) analysis plots of GeoMx spatial transcriptomic data from microregions (MRs) representing melanocytes and tumor cells, with colors indicating histopathological annotations (**A**), specimen stage, AJCC 8^th^ edition (**B**), and patient ID and sample set (**C**). Annotations are shown in **Supplementary Fig. S4A** and **S4B**. Each data point represents a single GeoMx MR. **D-H**. PC analysis plots of GeoMx spatial transcriptomic data from MRs representing melanocytes and tumor cells, colored by the enrichment scores for previously published melanoma-related and Hallmark gene signatures: *Joensson_Melanoma_Normal* (**D**)*, Landsberg_dedifferentiation_up* (**E**)*, Tirosh_MITF* (**F**)*, Joensson_Melanoma_Pigmentation* (**G**)*, Gopalan_Mesenchymal_Early* (**H**). **I**. PC analysis plots of GeoMx spatial transcriptomic data from melanocytes and tumor cells, colored by the solar elastosis score (specimen-level annotation). **J.** CyCIF images of specimens MEL82, MEL70, MEL74, and MEL72 stained with a subset of tumor, immune, and stromal markers: DNA (blue), and melanocyte/tumor (SOX10: red, NGFR: yellow, MART1: cyan), CD8^+^ T cells (CD8: orange), endothelial cells (CD31: white), CD206^+^ macrophages (CD206: green). Top row shows low magnification view, with rectangles corresponding to the ROIs magnified in the middle and bottom rows. Scale bars, 1 mm and 50 μm.

The enrichment of embryonic/neural crest gene sets to the lower right of the PCA landscape was most evident in three deeply invasive specimens (MEL70, MEL72, and MEL74; solid outline in **Fig. 3F,H**) which also exhibited uniformly high NGFR staining in the tumor compartment. Tumor cells arranged in these specimens had a morphology consistent with “vascular mimicry”, a histological feature associated with malignancy (**Fig. 3J**)^70^ and expressed the highest levels of *NGFR* and the lowest levels of *MLANA* and *MITF* of all specimens (**Supplementary Fig. S5E-G**). We identified one specimen (MEL82; **Fig. 3H** dashed outline) in which NGFR^+^ cells were intermixed with NGFR^-^ tumor cells that expressed SOX10, MART1, and/or SOX9 (**Fig. 3J, Supplementary Fig. S5H**). However, NGFR^+^ cells (and embryonic gene signatures in MRs) were rare or undetectable in earlier progression stages, showing that, in our cohort, an embryonic (neural crest-like) state is unlikely to be involved in melanoma initiation in most tumors.

Reconciling systematic variation in MR biology across the entire cohort with the intratumoral heterogeneity described in **Fig. 1 & 2** implies that MRs from a single specimen must differ from each other. We found that this was the case: for many specimens, MRs from that specimen were as different from one another as from MRs from all other tumors (note that this analysis used Set A tumors to minimize the impact of the uniformly NGFR-high tumors in Set B; **Fig. 4A, Supplementary Fig. S8A,B**). For example, for MEL18 (a stage IIA melanoma), some MRs with RGP and VGP regions mapped to the left in PC1 (MEL18_015 and _016) and others to the right (MEL18_026 and _027; **Fig. 4B**). The former MRs corresponded to cells growing along the follicular epithelium and invading beyond the follicular adventitia into the surrounding dermis, whereas the latter MRs were located in the dermal invasive component without hair follicle involvement (**Fig. 4B**). In the case of MEL25 (a stage IA melanoma), MRs corresponding to different invasion depths were spread along PC1 with MRs corresponding to the dermal invasive regions (regions MEL25_59 and _65) distinct from the intraepidermal component with superficial invasion (left regions MEL25_61 and MEL25_62, **Supplementary Fig. S8C,D**). Differences in transcriptional signatures between MEL18 and MEL25 MRs were readily discernable in the corresponding CyCIF images, for example partial loss of MART1 in MRs mapped to the left along PC1 and retention of MART1 staining in MRs mapped to the right (**Fig. 4A-C** and **Supplementary Fig. S8C-E**). These differences were not simply related to the distance between an MR and the dermal-epidermal junction (DEJ), an important consideration since the cellular composition of the skin changes dramatically from epidermis to dermis (**Supplementary Fig. S4D**).

**Figure 4.**
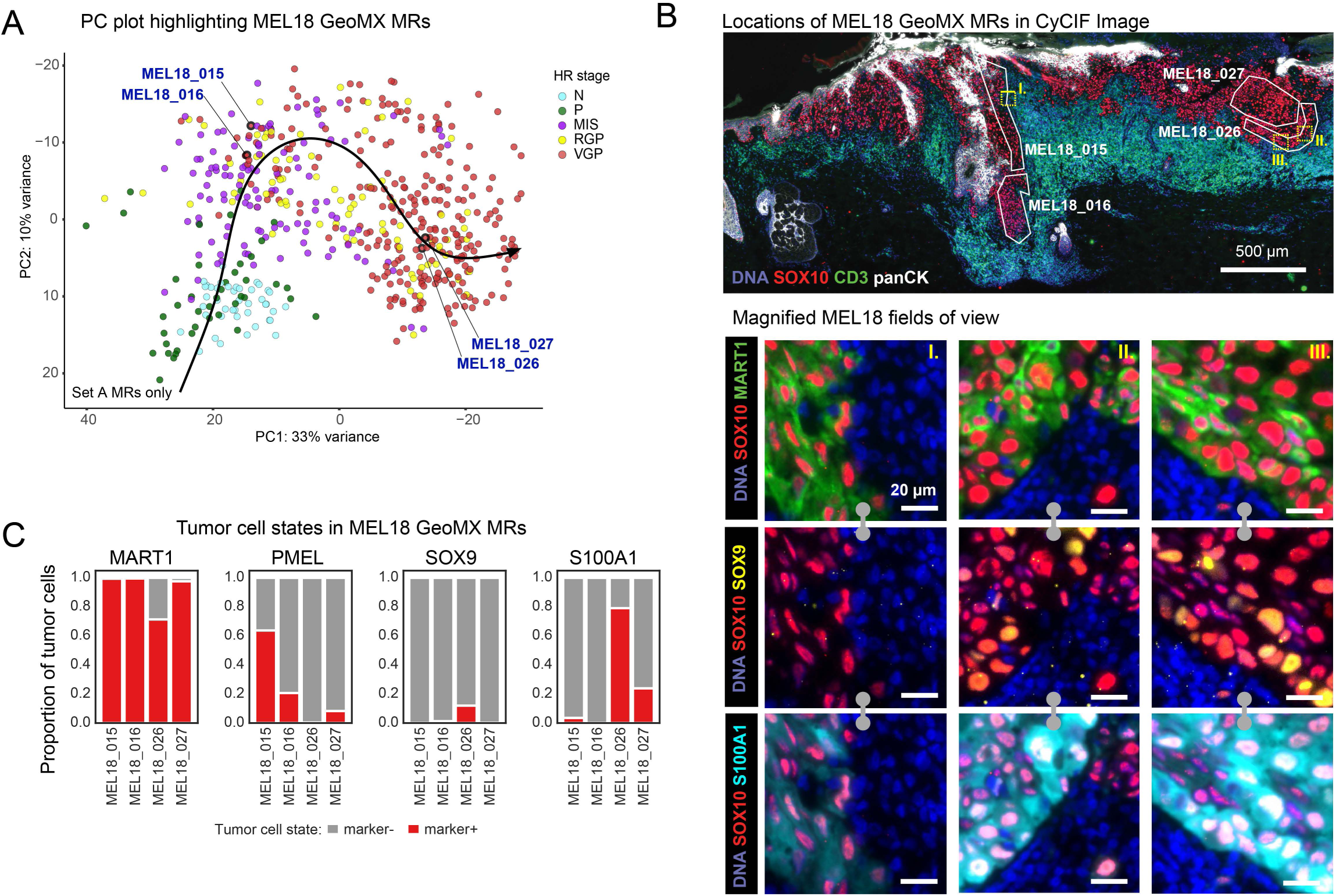
Tumor state heterogeneity within GeoMx MRs. **A.** PC plot of all GeoMx spatial transcriptomic data from microregions (MRs) representing melanocytes and tumor cells in sample Set A, colored by histological annotations. Each data point represents a single GeoMx MR. Four exemplary MRs from specimen MEL18 are highlighted (MRs MEL18_015, MEL18_016, MEL18_027, MEL18_026). **B.** Top: CyCIF image of MEL18 labeled with the spatial locations of selected VGP GeoMx microregions (white polygons). MRs are labeled with MR ID, indicating how the regions correspond to the PC plot in panel C. Lower: Magnified ROIs from top panel showing various tumor cell state markers, including SOX10 (red), MART1 (green), SOX9 (yellow). Scale bars, 500 μm and 20 μm. **C.** The proportion of tumor cells positive for MART1, PMEL, SOX9 and S100A1 within selected VGP GeoMx microregions from specimen MEL18.

These data show that the mesoscale cellular neighborhoods corresponding to MRs (200-600 cells or 2-10% of the disease-affected area of a specimen) represent a biologically significant spatial scale for disease-relevant differences in the tumor compartment of primary melanomas (note that these cell numbers represent a 2D slice; a symmetrical 3D MR might therefore be expected to comprise ∼ 5 x 10^3^ cells). The fact that MR biology can be well described by changes in expression of MITF-regulated and embryonic gene signatures, both of which were first identified in late-stage melanoma, suggests that key features of metastatic disease emerge early in progression. However, these tumor states are not generally present in a pure form either in a single specimen or within a single histological domain (as defined by progression stage) but instead were intermixed at the level of local neighborhoods. Moreover, the continuity of tumor states in PCA space and evidence of continuous variation in protein expression in CyCIF images also suggests that the transition from a normal to an invasive state involves continuous rather than discrete changes in gene expression.

### Progression programs in early-stage melanoma

As a complementary approach to analyzing gene expression across MRs we used machine learning based on self-organizing maps (SOMs); this approach projects high-dimensional gene expression onto a two-dimensional grid.^47^ We concentrated on earlier stage disease (Set A) and modeled the two batches of GeoMx signatures separately since SOMs were more sensitive to batch effects than PCA analysis (n = 299 MRs for Set A1 and n = 239 for Set A2); this yielded 900 metagenes (sets of genes with coordinated expression profiles). We mapped the metagenes onto a two-dimensional representation of the SOM (the SOM portrait) based on their expression pattern and decomposed portraits for each local progression stage. This revealed a well-organized progression series: MRs associated with normal and precursor stages projected onto the lower right corner of the portrait, RGP and VGP stages projected onto the upper left corner, and MIS stages exhibited intermediate projection (**Fig. 5A** and **Supplementary Fig. S9A,B**). Thus, the SOM had learned the progression axis from the original data in a fully unsupervised manner, again demonstrating that inter- and intra-tumoral heterogeneity were not random but were associated with an underlying spatial organization.

**Figure 5.**
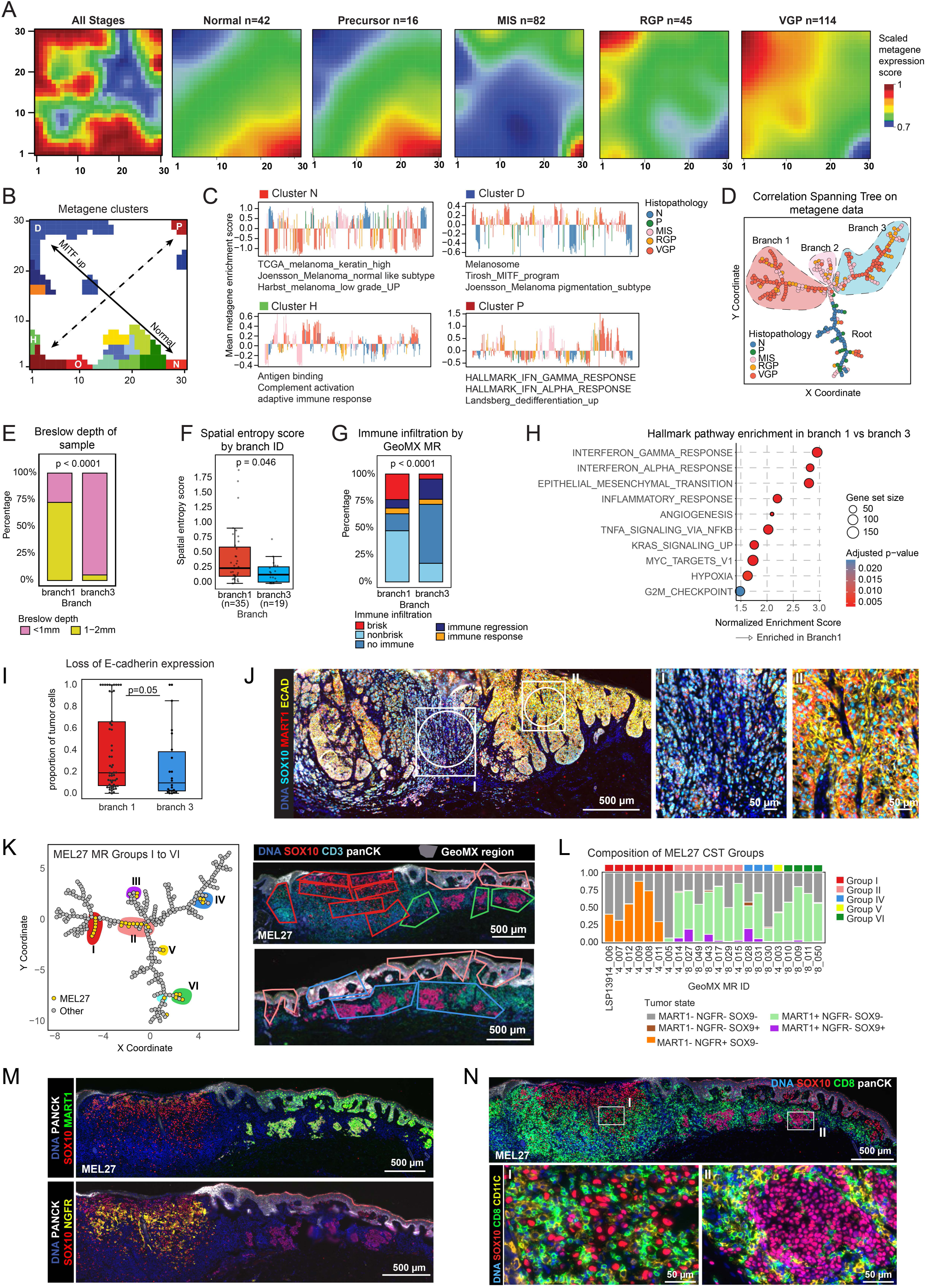
Developmental trajectory from normal melanocytes to invasive melanoma. **A.** Self-organizing map (SOM) portrait of GeoMx data (Set A1, n = 299 from n = 28 patients), showing the global expression of 900 metagenes across profiled samples (left). Each square within the 30x30 grid represents an unique metagene. SOM portraits for local progression stages show the global expression of 900 metagenes in microregions (MRs) grouped by histopathology. The expression levels of each metagene are indicated with a color (scaled metagene expression score) as compared to the average expression of all metagenes. **B.** SOM clusters A-Q and their mapping to the SOM portrait. Unique SOM clusters are indicated with unique colors. Spots N, D, H, P and O are highlighted, and the rest are shown in **Supplementary Fig. S9F**. **C.** Mean expression profiles of the defined SOM clusters N, D, H and P within microregions from various progression stage categories. The Y axis represents the mean expression profile of the SOM cluster signatures of each microregion. Individual MRs are shown in the X axis. The pathways listed below represent the most enriched pathways within each SOM cluster. **D.** Minimum spanning tree analysis showing the correlation of profiled GeoMx MRs. The datapoints (colored circles) represent individual MRs. The correlation spanning tree (CST) is annotated by the local progression stage for each MR and the branch identity (Branches 1, 2 and 3). **E.** The percentage of MRs in Branches 1 and 3 belonging to samples with Breslow depth <1mm and 1-2 mm. P-value was calculated based on Fisher’s exact test. **F.** Spatial entropy scores calculated for the corresponding tumor regions in the adjacent tissue section for Branch 1 and 3 MRs. Each data point represents a spatial entropy score calculated for individual GeoMx MRs, and the samples are grouped by the CST branch identity. P-value was calculated based on Wilcoxon Rank Sum test. **G.** The proportion of annotations for immune responses (brisk TIL, non-brisk TIL, no immune, immune regression, immune response) within Branch 1 and 3 MRs. P-value was calculated based on Fisher’s exact test. **H.** Gene-set enrichment analysis (GSEA) of Hallmark gene set (pathways) significantly enriched in Branch 1 and Branch 3 microregions. Dot size indicates the size of the gene set and color indicates the adjusted *p* value. **I.** The proportion of tumor cells staining positive for E-cadherin of all tumor cells in Branch 1 and Branch 3 microregions. Each data point represents the mean proportion of marker+ cells within a single GeoMx MR, and the samples are grouped by the CST branch identity. **J.** CyCIF image of specimen MEL54 stained for DNA (blue), SOX10 (cyan), MART1 (red), E-cadherin (yellow), highlighting two GeoMx microregions from Branch 1. Scale bars, 500 μm and 50 μm. **K.** Left: CST highlighting microregions from specimen MEL27 (yellow dots). The grey datapoints represent individual MRs from other patients. Groups of microregions are encircled with different colors and annotated with roman numerals (I-VI). A subset of these microregions is highlighted in the CyCIF images on the right. **L.** The proportion of various tumor cell states within selected VGP GeoMx microregions from specimen MEL27. Cluster ID indicates the grouping of MRs shown in the correlation spanning tree in panel K. **M.** CyCIF image of specimen MEL27 stained for DNA (blue), SOX10 (red), panCK (white), MART1 (green; top row), and NGFR (yellow; bottom row). **N.** Top: CyCIF image of specimen MEL27 stained for DNA (blue), SOX10 (red), panCK (white) and CD8 (green). Bottom: magnified regions indicated with I and II are stained for DNA (blue), SOX10 (red), CD8 (green) and CD11C (yellow).

To identify transcriptional features contributing to the SOM portrait, metagenes were clustered based on a min-max scaled threshold of 0.95, corresponding approximately to the top 5% of expression value in each metagene profile; this yielded 17 clusters of co-expressed metagenes. These clusters represent key discriminating features in the data and as expected, mapped primarily to the edges of the SOM portrait (Clusters A-Q; **Fig. 5B,C**; **Supplementary Fig. S9C,D**). Cluster N, which existed in the lower right region of the portrait associated with normal and precursor MRs, was enriched in gene expression signatures such as “*Joensson_Melanoma_normal like subtype”* (see **Fig. 2** and **3**). Cluster D in the upper left of the portrait, the location of RGP and VGP MRs, was enriched in signatures of MITF upregulation (e.g., “*Tirosh_MITF_program*, **Fig. 5C**). Moreover, when the clusters were projected back to the PC landscape, enrichment scores for clusters N and D corresponded well to the progression trajectory along PC1 (**Supplementary Fig. S9E**). The other diagonal of the SOM portrait (dotted line in **Fig. 5B**) was organized by differences in immune and inflammatory signatures, with IFNγ and IFNα signatures in the upper right (cluster P) and other immune features in the lower left (cluster H; **Fig. 5B,C**; see **Supplementary Fig. S9F** for data on all clusters). Dedifferentiation signatures were not a major organizing principle it this particular SOM organization (e.g. *Landsberg_dedifferentiation_up*), probably because dataset A1 was enriched in earlier stage specimens with fewer NGFR^+^ tumors (the portion of the PCA plot depicted in **Supplementary Fig. S4A,C**).

To identify the information learned by the SOM we calculated the pairwise correlation among metagenes (Pearson’s correlation coefficient, **Supplementary Fig. S9G, Supplementary Fig. S10A**) and then generated a correlation spanning tree (CST, **Fig. 5D, Supplementary Fig. S10B**). A CST represents a cluster map as a partitioned tree, a representation that is easy to parse but does not reduce the fidelity of clustering.^71^ We interpreted the resulting CST as having a root and three branches, with the root comprising primarily normal and precursor regions (**Fig. 5D** microregions in blue and green), branch 2 comprising MIS regions, and branches 1 and 3 comprising VGP and RGP. Comparing the invasive branches, we found that branch 1 was enriched in MRs from tumors with Breslow depth >1 mm (**Fig. 5E**) and corresponded to clusters D and P of the SOM portrait (**Supplementary Fig. S10C**). MRs in branch 1 had higher spatial entropy (**Fig. 5F** and **Supplementary Fig. S10D**) and greater immune-association than MRs from the thinner and more melanocytic (MITF-high) MRs in branch 3 (**Fig. 5G**). Enrichment analysis confirmed that branch 1 was significantly enriched in IFNγ, IFNα, and TNFα responses, and in epithelial-mesenchymal transition (EMT signatures, **Fig. 5H**). To confirm this finding, we compared E-cadherin staining in branch 1 and 3 MRs. E-cadherin is a cell adhesion protein and tumor suppressor, whose loss is a hallmark of EMT commonly observed when melanocytes lose their physical and regulatory links to keratinocytes.^72^ We observed a significantly higher proportion of E-cadherin negative cells in branch 1 than branch 3 MRs (**Fig. 5I,J**); overall, E-cadherin intensity levels were positively correlated with MART1 levels (**Supplementary Fig. S10E**). Thus, inflammatory and EMT signatures that were not well resolved by PCA analysis were revealed by the SOM to comprise two (and possibly more) distinct trajectories.

When MRs making up the CST were labeled by specimen identity, most specimens were found to contain MRs that mapped to multiple branches (**Supplementary Fig. S10F**), consistent with the high intratumoral heterogeneity observed visually and by PCA analysis (**Fig. 3**). For example, stage IB specimen MEL27 formed 6 groups based on their position in the CST (labelled I to VI; **Fig. 5K, Supplementary Fig. S10G**). MRs in group I were enriched in MART1^-^ NGFR^+^ tumor cells (and a brisk CD8+ T cell infiltrate was present), while group II MRs contained MART1^+^ NGFR^-^ cells (**Fig. 5L-N**). MRs in groups II, IV and VI represented the superficial or intraepidermal component of invasive melanoma (**Fig. 5K**). In contrast, MEL14 was an example of a tumor in which all tumor MRs clustered closely together in Branch 1 (**Supplementary Fig. S11A**). These MRs were also similar to each other with respect to tumor cell states, with 92% of tumor cells being MART1^+^ PMEL^+^ (**Supplementary Fig. S11B,C**). Unsupervised analysis therefore confirms our previous conclusion that MRs from a single tumor can be as variable as MRs from the entire cohort but that high intra-tumoral heterogeneity is not an obligatory feature of primary melanoma.

### Spatial features of tumor microenvironment (TME) in melanoma dedifferentiation

To identify spatial features distinguishing CST branches 1 and 3, we used K-means clustering (*k* = 25) to generate recurrent spatial cellular neighborhoods (RCNs), which separate intermixed cell types into recurrent communities based on local proximity and relative enrichment (**Fig. 6A,B**). The RCNs were then clustered into groups (RCNGs; n = 11) that corresponded to (i) components of the skin such as epidermis, stroma, or vasculature, (ii) immune domains of different types, (iii) tumor domains, and the (iv) tumor-stromal interface (**Fig. 6B-D**). The abundance of epidermis RCNGs (RCNG4) decreased by 10-fold from precursor to VGP while T cell and tumor RCNGs (RCNGs 5 and 1) increased by ∼5 - and 25-fold, respectively. In contrast, RCNGs for myeloid/macrophage (RCNG7) and vascular (RCNG11) varied less than 3-fold (**Fig. 6E**). However, the proportions of RCNG7 and RCNG11 were significantly more abundant in the MRs in branch 1 than 3 (**Fig. 6F,G**, **Supplementary Fig. S12A,B**). RCNG7 corresponded to a myeloid-rich compartment located either near the invasive front or instead within the dermis as patches alternating with T cells in neighborhood RCNG5. RCNG11 corresponded to blood vessels, visible in **Fig. 6D** in both longitudinal and transverse aspects. To control the possibility that depth-dependent differences in skin composition might confound this analysis, we tested for differences in RCN prevalence with Breslow depth; however, no significant associations were observed (**Supplementary Fig. S12C**). Enrichment of branch 1 MRs for proximity to RCNG11 (myeloid cells) but not RCNG5 (T cells) implies that T cells and myeloid cells were not highly intermixed in most specimens and could therefore differentially pattern the TME. Consistent with this observation, proximity analysis showed that CD8^+^ T cells were on average the nearest neighbor immune cell type for other CD8^+^ T cells and macrophages were the closest neighbors to other macrophages (**Supplementary Fig. S12D**).

**Figure 6.**
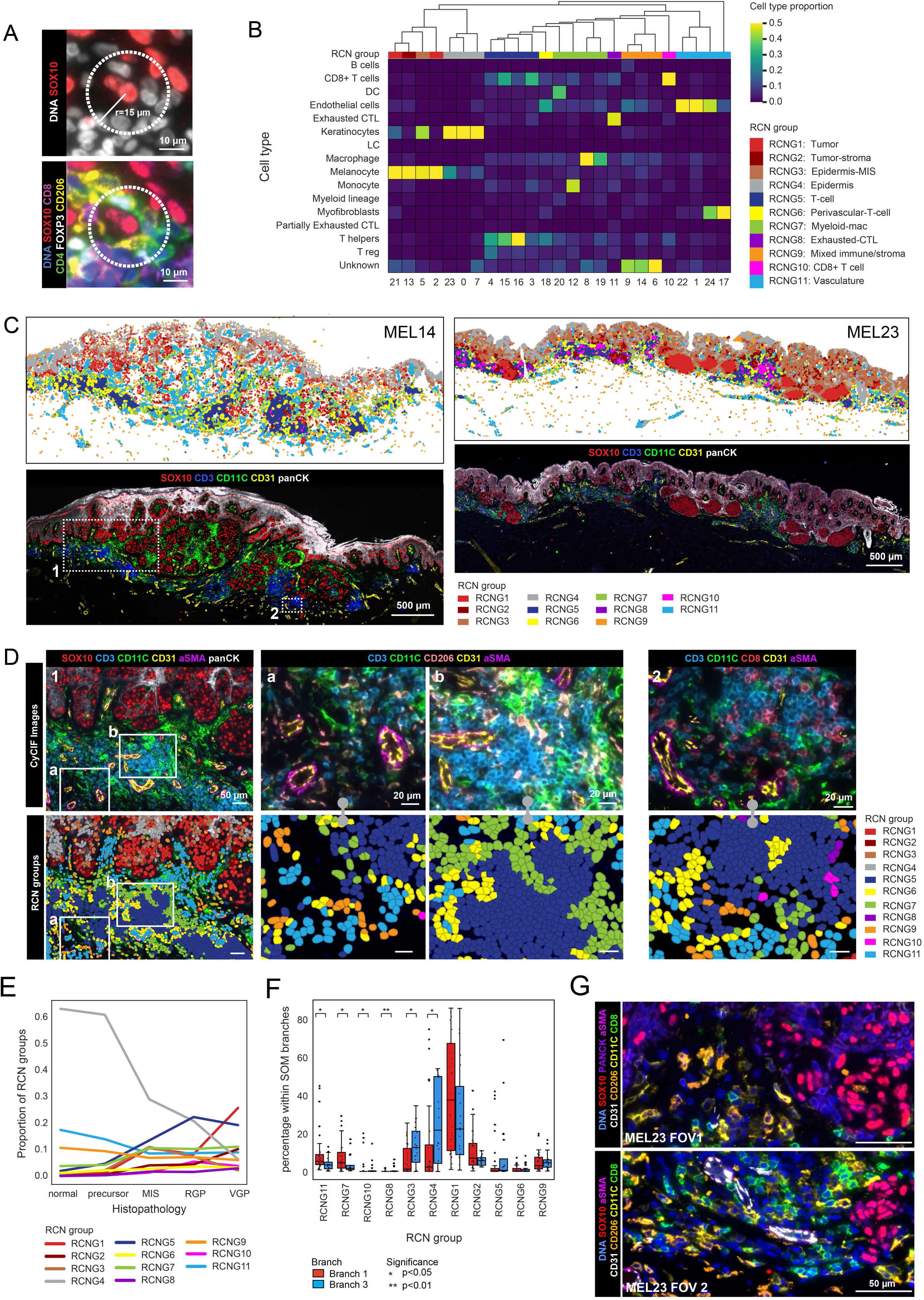
Spatial features of tumor microenvironment (TME) in melanoma dedifferentiation. **A.** CyCIF image of a field of view in specimen MEL18 showing an example of a cellular neighborhood with a 15 μm (50 px) radius, stained for DNA (blue), SOX10 (red; left). The corresponding region (right) stained for DNA (blue), SOX10 (red), CD8 (purple), CD4 (green), FOXP3 (white), CD206 (yellow). Scale bars, 10 μm. **B.** A heatmap showing the mean proportion of cell types within Recurrent Cellular Neighborhoods (k=25) and their grouping into 11 meta clusters (RCN groups: RCNG). **C.** Top: Scatter plots showing the RCN annotations for each cell within selected fields of view for samples MEL14 (left) and MEL23 (right). Each datapoint represents a cell centroid in the corresponding CyCIF image. The corresponding CyCIF image is shown below for both tumors, stained for SOX10 (red), CD3 (blue), CD11C (green), panCK (white), CD31 (yellow). Scale bars, 500 μm. **D.** Magnified regions 1 and 2 from specimen MEL14 shown in panel C. Top left: Region 1 stained for SOX10 (red), CD3 (blue), CD11C (green), panCK (white), CD31 (yellow), aSMA (magenta). Bottom left: Region 1 with RNCG annotations for each single cell. Middle: magnified insets a and b from Region 1 stained for CD3 (blue), CD11C (green), CD206 (orange), CD31 (yellow), aSMA (magenta), and their corresponding RCNG annotations. Top right: Region 2 stained for CD3 (blue), CD11C (green), CD8 (red), CD31 (yellow), and aSMA (magenta). Bottom right: Region 2 with RNCG annotations for each single cell. Scale bars, 50 μm and 20 μm. **E.** The proportion of RCN groups (RCNG1-11) within local progression stage HRs (normal, precursor, MIS, RGP, VGP). **F.** RCNG identities of single cells belonging to Branch 1 and Branch 3 microregions. Box plots show the proportion of cells within Branch 1 and Branch 3 belonging to different RCNGs in each CyCIF sample. Each datapoint represents the mean percentage of RCNGs within each sample, and the samples are grouped by their correlation spanning tree identity (Branch 1 vs Branch 2). *, P < 0.05; **, P < 0.01. **G.** Magnified fields of view from specimen MEL23 (CyCIF image shown in **Supplementary Fig. S12B**). Staining for DNA (blue), SOX10 (red), panCK (purple; top panel only), aSMA (magenta), CD31 (white), CD206 (orange), CD11C (yellow), and CD8 (green) are shown. Scale bars 50 μm.

Moreover, branch 1 MRs were found (by CyCIF) to contain a higher proportion of CD163^+^ and/or CD206^+^ macrophages and CD31^+^ endothelial cells than branch 3 MRs (**Supplementary Fig. S13A**). We conclude that proximity to myeloid cells was one factor distinguishing branch 1 & 3 cells and that proximity to a vascular/perivasculature environment, an established negative prognostic feature in primary melanoma^73^, is another. Thus, it appears that mesoscale architectural features such as blood vessels and clustering of immune cells (myeloid cells in particular) represent substantial contributors to intratumoral heterogeneity.

### Quantifying tumor cell state heterogeneity within an MR using spatial entropy

When CyCIF images from individual MRs were examined closely using high dimensional CyCIF it was apparent that tumor cells were not as homogeneous as we had initially postulated when designing the GeoMX sampling strategy (**Fig. 2B, 3J, 4B**). For example, both superficial and deep MRs from MEL18 and MEL25 contained neighboring tumor cells expressing SOX9, SOX10 and MART1 in different combinations and intensities, consistent with frequent changes in molecular programs controlling these melanocyte markers (**Fig. 4B,C, Supplementary Fig. S8D,E**). We quantified intermixing using local Shannon entropy, with neighbors defined by Delaunay triangulation on tumor cells (ignoring stromal or immune cells), and cell states as defined in **Fig. 2A**. A low entropy score corresponded to a region of tissue in which tumor cells were similar in type, while a high entropy score corresponded to a spatially intermixed state. Using this approach, we found that some specimens (e.g., MEL18) exhibited low overall entropy whereas others (e.g., MEL25) exhibited high entropy (**Fig. 7A**). However, any single specimen could have both high and low local entropy domains, as evidenced by variability in entropy maps at both the whole-specimen and local levels (**Fig. 7A,B**).

**Figure 7.**
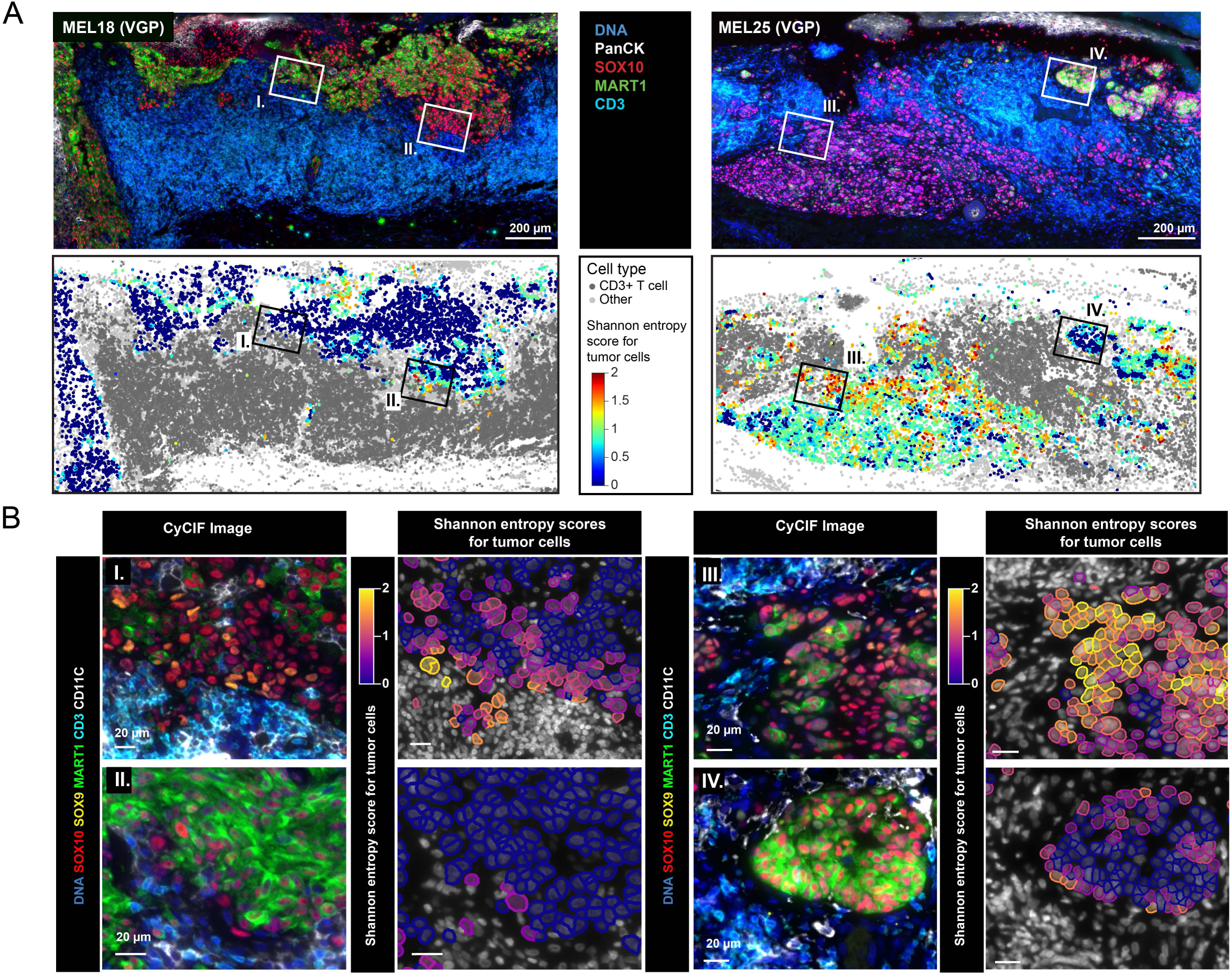
Quantifying tumor cell state heterogeneity with spatial entropy. **A.** Top: CyCIF image of specimens MEL18 (top left) and MEL25 (top right) stained for DNA (blue), epidermis (panCK; white), melanocytes (SOX10; red), tumor (MART1; green) and T cells (CD3; cyan). Bottom: Scatter plots show the local neighborhood Shannon entropy values of each SOX10^+^ tumor cell in specimens MEL18 and MEL25, which are mapped back to the corresponding tissue location using the X,Y coordinates of the cell centroids. Each data point represents a cell centroid. The centroids for CD3+ T cells and other non-tumor cells are shown with dark and light grey dots, respectively. The boxes labeled as I-IV represent the regions magnified in **B**. Scale bars 200 μm. **B.** Magnified CyCIF ROIs from **A**, stained for DNA (blue), SOX10 (red), MART1 (green), SOX9 (yellow), CD3 (cyan), CD11c (white). The adjacent plots showing segmented tumor cells show the local neighborhood Shannon entropy values calculated for each SOX10+ tumor cell within the magnified regions. Scale bars 20 μm.

To identify factors associated with variation in entropy within MRs, we examined MR gene expression profiles and local histology (the spatial entropy used for calculation of MR entropy differed subtly from the single cell entropy described above; see Methods; **Fig. 8A,B**). We observed that, on average, spatial entropy increased significantly with histological stage (e.g., from MIS to RGP or MIS to VGP). NGFR^+^ locally advanced specimens with Breslow depth >4mm in set B were an outlier in this analysis and had the lowest entropy of all specimens in the cohort (**Supplementary Fig. S14D-F**); whether this represents a general evolution of tumor state to greater homogeneity or is a feature of a minority of specimens is not yet known. When mean entropy values for all HRs in a specimen were compared to Breslow depth for that tumor, we found that the average spatial entropy value was constant from thinner to thicker specimens, but the variance increased significantly (as measured by Levene’s test; **Fig. 8B**). Since Breslow depth is a specimen-level measurement, we also divided specimens into bands perpendicular to the direction of invasion. In this case, we did not observe a consistent or significant association between local depth of invasion and spatial entropy (**Supplementary Fig. S14A-C**): in many specimens the spatial entropy value was constant with depth, whereas in others it decreased (e.g., MEL18) or increased with depth (MEL58; **Supplementary Fig. S14B**). Thus, it was not the position of cells within a single tumor, as approximated by depth of invasion, that determined the degree of entropy, but rather the overall thickness of the specimen (the Breslow depth), which is a measure of progression. These data demonstrate that the degree of intermixing of tumor cell lineages (the spatial entropy) increases from MIS to VGP stages and the variability in the entropy value also increases as specimens get thicker overall (**Fig. 8B**). These data show that seemingly random fluctuation in state at the length scale of single cells is overlaid in some specimens on changes in mean gene expression at the level of MRs.

**Figure 8.**
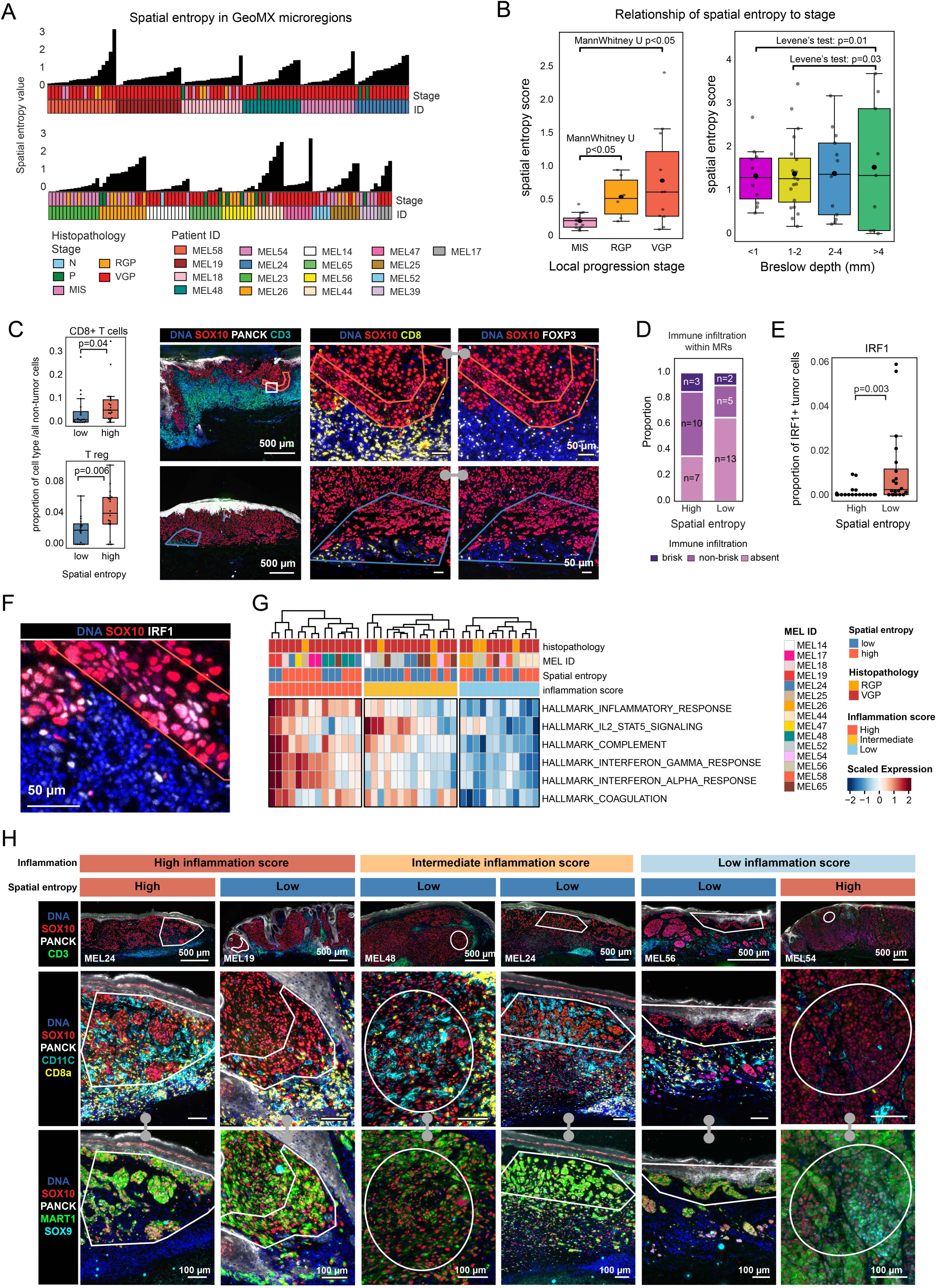
Spatial entropy and inflammation. **A.** CoMut plot of the spatial entropy scores for each GeoMx microregion (MRs), calculated using CyCIF data of the adjacent tissue section. Each MR is annotated for histopathology (N, P, MIS, RGP, VGP) and patient ID. **B.** Spatial entropy scores calculated by local progression stages (left) and Breslow depth (right; for RGP and VGP regions only). Each data point represents the spatial entropy score calculated for a histologic region in a single sample. Box represents the first and third interquartiles of the data and whiskers extend to show the rest of the distribution except for points that are determined to be outliers. The p-value of local progression stage comparison was calculated using MannWhitney U test. The p-value of Breslow depth comparison was calculated based on Levene’s test. **C.** Left: The proportion of CD8^+^ T cells (top) and T regs (bottom) out of all non-tumor cells within high or low entropy microregions. Each data point represents a mean cell type proportion within a single GeoMx MR, and the samples are grouped by the MR spatial entropy scores. Box represents the first and third interquartiles of the data, whiskers extend to show the rest of the distribution except for points that were determined to be outliers. Right: CyCIF images of samples MEL18 (top), and MEL65 (bottom), stained for DNA (blue), panCK (white) and SOX10 (red), CD3 (cyan). The magnified box in specimen MEL18 indicates a microregion with high spatial entropy, and in specimen MEL65 indicates microregion with low spatial entropy. These regions are magnified next to the whole-slide images and stained for DNA (blue), SOX10 (red), CD8 (yellow) and FOXP3 (white). Scale bars, 500 μm and 50 μm. **D.** Stacked bar plot showing the distribution of annotations for immune cell infiltration as evaluated by CD45 staining for GeoMx MRs. The proportion of various immune cell infiltration patterns is shown for MRs with high and low spatial entropy. The p-value was calculated using Chi-square test. **E.** Boxplot showing the proportion of IRF1^+^ tumor cells within high and low entropy MRs. Each data point represents the mean proportion of marker+ cells within a single GeoMx MR, and the samples are grouped by the MR spatial entropy scores. The p-value was calculated using MannWhitney U test. **F.** Magnified field of view of a high spatial entropy MR shown in panel **C** (specimen MEL18), stained for DNA (blue), IRF1 (white) and SOX10 (red). Scale bar, 50 μm. **G.** The enrichment scores of Hallmark gene set (pathways) for spatial entropy MRs (n = 40), annotated by histopathology (RGP, VGP), spatial entropy score grouping (high, low), patient ID, and inflammation score (high, intermediate, low). **H.** Top row from left to right: CyCIF images of samples MEL24, MEL19, MEL48, MEL24, MEL56 and MEL54, stained for DNA (blue), panCK (white), SOX10 (red), and CD3 (green). White regions indicate individual exemplary GeoMx MRs annotated for inflammation score and spatial entropy (annotations shown on top of each column). These regions are magnified below the low power fields of view and stained for DNA (blue), SOX10 (red), MART1 (green), CD3 (yellow) and CD11C (cyan; middle row); DNA (blue), SOX10 (red), SOX9 (cyan), and MART1 (green; bottom row). Scale bars, 500 μm and 100 μm.

Host immune response has previously been shown to promote lineage plasticity in advanced melanoma^26^ and it has been suggested that IFNγ is a primary driver of this plasticity.^74^ We therefore asked whether spatial entropy at the level of MRs was associated with proximity to an inflammatory environment. First, we used CyCIF to determine the immune cell composition of MRs having high (n=20) and low (n=20) spatial entropy in the tumor compartment. We found that high entropy regions contained significantly higher proportions of CD8^+^ T cells and CD4^+^ FOXP3^+^ regulatory T cells than low entropy regions (**Fig. 8C**, **Supplementary Fig. S14G**) and were qualitatively more immune infiltrated, as judged visually using criteria similar to how brisk TIL infiltrates are scored clinically^75^ (**Fig. 8D**, **Supplementary Fig. S14H**).

Tumor cells in high entropy MRs were also significantly more likely to have nuclear IRF1 (interferon regulatory transcription factor 1), suggesting that they were exposed to IFNγ (**Fig. 8E,F**). Based on gene set enrichment (ssGSEA), high entropy MRs were associated with inflammatory signatures, such as responses to IFNα and IFNγ, classic inflammatory cytokines, the complement pathway, a component of innate immunity, and coagulation (note that the coagulation and complement pathways also contain cytokines and chemokines or their receptors;^76^ **Fig. 8G,H**). As an alternative approach we examined the local environment of tumor cells that had lost MART1 expression, which is a univariate measure of tumor cell dedifferentiation and among the most variable of all tumor markers. Using proximity analysis to compare cells within a 15 μm radius of either SOX10^+^ MART1^-^ or SOX10^+^ MART1^+^ tumor cells, we observed a significant association between loss of MART1 expression and CD8^+^ T cells, CD4^+^ Treg cells, macrophages, and endothelial cells (**Supplementary Fig. S13B,C**). We conclude that high entropy in the tumor compartment, and its correlate, tumor cell plasticity, as well as loss of MART1, and its correlate, tumor cell differentiation, are both significantly associated with proximity to immune cells and responsiveness to inflammatory cytokines.

However, at the level of individual MRs, spatial entropy and local immune reaction were not obligatorily linked: some MRs with high spatial entropy did not exhibit upregulation of inflammatory gene signatures and were not proximate to immune cell populations; the converse was also true (**Fig. 8G,H).** This lack of one-to-one association was also evident in maps of Shannon entropy at single cell resolution: some regions with abundant T cells in close proximity to tumor cells were homogenous and exhibited low Shannon entropy values at single-cell level (**Fig. 7A,B**). Thus, while inflammation can cause spatial entropy, other (unmeasured factors) must be additional contributors.

## DISCUSSION

The high inter- and intra-tumoral heterogeneity observed in primary cutaneous melanoma makes it challenging to study molecular mechanisms promoting tumor initiation, progression, and invasion and greatly complicates diagnosis and risk stratification. Understanding the origins of this heterogeneity, and determining what it means clinically, is likely to be important in improving the treatment of melanoma, particularly when the disease is still curable by surgery (or in high-risk cases, in combination with neoadjuvant therapy)^77^ while minimizing overdiagnosis.^6^ In this paper, we establish that inter- and intra-tumoral heterogeneity in primary melanomas is not random, but is instead organized with respect to two fundamental axes that specify melanocyte fate: the activity of the MITF transcriptional regulator and expression of neural crest genes associated with a dedifferentiated phenotype. The strength of MITF signatures and the expression of MITF target genes such as *MLANA* (MART1) are intermediate in normal skin and precursor regions, elevated in many RGP and VGP domains (which also over-expressed pigmentation genes), and lowest in invasive VGP domains. This finding is consistent with the rheostat model of MITF inferred from scRNA-Seq of metastatic tumors and melanoma cell lines.^78^ The de-differentiated state has also primarily been characterized in metastatic tumors and involves elevated expression of genes associated with progenitor neural crest cells (and with NGFR protein expression)^29^. Across all specimens and histological domains in our dataset, the average trajectory of cell states in this MITF v. neural crest landscape was well correlated with local histopathology, demonstrating the existence of defined progression “pathway”. Many of the cell state changes observed along this pathway are described as therapy-induced^79^, but we find them to be common in early stage treatment-naïve cancers.

Melanoma cell states are determined not at the level of whole specimens or histological domains but rather locally at the level of MRs. Within a single tumor these MRs vary substantially relative to the average trajectory; thus late-stage tumors often have features characteristic of earlier progenitors and some early-stage tumors have MRs with expression programs characteristic of deep invasion. High variation in tumor cell state from one local region to the next in a single tumor is consistent with the well-established plasticity of melanoma cells^11,80^ and it raises the question: what factors drive plasticity and determine cell state? One possibility is diffusible factors and cell-cell contacts in the local microenvironment. Reconstruction of MR state spaces using SOMs and spatial statistics supports this idea and implicate proximity to immune cells and the resulting induction of inflammatory programs.^12^ This is consistent with evidence that plasticity^80^ and dedifferentiation^27^ can be induced by an inflammatory environment in cell lines. Proximity to perivascular spaces is a second factor implicated in fate specification, and this is consistent with the role of angiotropism as a negative prognostic feature in primary melanoma^73^ and metastatic disease.^81^ However, our data demonstrate that these are not obligatory relationships; for example, melanoma cells can remain in an early progression stage in the face of proximate immune environments, implying the existence of additional spatial regulators.^82^

A second non-exclusive possibility is that tumor cell state is subject to frequent stochastic fluctuation resulting in a more or less random distribution of states. This does not appear to be a dominant effect at the length scale of MRs, but within some MRs we observe high levels of cell-to-cell variation in the expression of melanocytic markers (quantified here by spatial entropy). Some MRs exhibit high entropy at these short length scales and others do not – and once again proximity to inflammatory environments is implicated. Tumor cell-intrinsic factors are likely to be involved in this type of patterning, based on evidence that some melanoma tumor clones (or induced states) are simply more plastic than others.^82^ Genetic instability and clonal evolution are a third possibility, but their spatial properties remain frustratingly difficult to determine in small primary tumors. Stage II specimens such as MEL82 with clear differences in protein expression from one side of a tumor to another are candidates for functionally polyclonal primary cancers.

### Emergence of embryonic cell states during invasion

Many models of tumorigenesis postulate that embryonic states resembling self-renewing stem cells arise early in disease^83^ and promote the development of a diversity of downstream states in response to pressure from immunosurveillance or therapy.^84^ However, we find that the dedifferentiated states readily detectable in the invasive components of later stage melanoma are very rarely present in MIS or stage I melanoma; this is true whether induction of neural crest signatures, NGFR protein expression, or expression of embryonic genes is used as a criterion. Interestingly, the least diverse specimens in our cohort were deeply invasive Stage IIb/IIc melanomas. These occupied the extremum of the PCA state landscape and were uniformly populated by NGFR-high tumor cells with a vascular mimicry phenotype. By multiple criteria, these cells were more dedifferentiated than the cutaneous component of Stage III (locally metastatic) tumors also in the dataset. Clinically, deep primary tumors are strongly associated with worse outcomes, with Stage IIB/C disease (primary > 4mm depth invasion) having worse prognosis than shallower Stage IIIA tumors in which the cancer has metastasized to lymph nodes.^4^ One possibility is that deep tumors undergo purifying selection as they progress. Alternatively, uniformly NGFR-high tumors might have been “born bad” and progressed rapidly to deep invasion. Analysis of more specimens stratified by recurrence will be necessary to distinguish these possibilities.

### Factors patterning the melanoma TME and tumor compartment

A picture emerges from this study of tumors strongly influenced by proximity to spatially patterned features of the stroma such as the immune system, basement membranes, and vasculature.^12^ For example, decomposition of tumor states using SOMs reveals systematic differences in tumor states adjacent to myeloid or T cell-enriched stromal neighborhoods with proximity to myeloid RCNs associated with a more aggressive and “mesenchymal” state.^85^ Blood vessels are spaced every 100-200 µm in the epidermis, generating micro domains that can be predominantly arteriolar, venular, or devoid of microvasculature.^86^ Dedifferentiation and loss of expression of cell adhesion proteins such as E-cadherin in melanoma cells is associated with proximity to the vasculature, consistent with the role of angiotropism and embryonic programs in promoting melanoma invasiveness and metastasis.^81^ Additional stromal features we have not yet assessed include proximity to nociceptor neurons, which are known to influence immunosurveillance,^87^ and follicles, which may restrict melanoma spread.^88^

None of the significant associations we detect between the TME and tumor cell state are absolute: for example, tumor cell neighborhoods can be dedifferentiated and plastic without evidence of proximity to immune cells and the converse is also true. This suggests that many regulatory interactions and their directionality remain to be identified. Moreover, there is no reason to believe that we have optimally sized the MRs in this study. Although identified based on local morphology, MRs still contained cells of multiple types. A more sophisticated approach to MR definition, the addition of more features of the normal skin, 3D maps of nerves and the vasculature, and more refined transcriptional profiling promise to reveal additional factors patterning primary melanomas.

### Diagnostic implications

Better prognostication of primary melanoma is urgently needed to discriminate low-risk tumors that do not require aggressive treatment or frequent surveillance from those requiring systemic therapy and close monitoring.^89^ Use of molecular test for prognostication remains however controversial^90^, and American Association of Dermatology guidelines do not recommend molecular testing (outside of clinical trials) due to insufficient evidence of efficacy.^91^ The current study raises four questions impinging on this controversy and prognostication more generally. First, does progression of a tumor along the MITF v. neural crest landscape correspond to risk of metastasis or recurrence and if so, how might these states best be measured in clinical specimens? Second, does a tumor that is largely at an early progression state but has some MRs with late-stage properties represent a high-risk tumor? Third, is high intratumoral variation (high cell state entropy) itself a risk feature? Fourth, to what extent are any of these properties correlated with or explanatory of established diagnostic features such as Breslow depth? It is noteworthy that prognostic tests currently being developed or marketed rely primarily on bulk gene expression profiling (e.g. CP-GEP^92^, Skyline DX®^93^, DecisionDX®^94^ etc.) that inevitably obscures intratumoral heterogeneity. If a risk feature is recessive (e.g., a reduction in gene expression in an MR) then it may be very difficult to reliably detect using bulk assays. Thus, spatial approaches that build on current dermatopathology practice such as multiplexed imaging combined with machine learning on H&E images^95^ may represent a superior approach.

## ACKNOWLEDGEMENTS

This work was supported by the Ludwig Center at Harvard an ASPIRE Award from The Mark Foundation for Cancer Research, and NCI grants U2C-CA233262 to PKS, SS; NCI grants R00-CA256497 to AJN and R50-CA252138 to ZM. TV is supported by a Research Scholar Grant, PF-24-1316850-01-CD, from the American Cancer Society. Histopathology support was provided by P30-CA06516. SS is supported by the BWH President’s Scholars Award. We thank Kerrie Marie and Vishaka Gopalan for their help and expertise in this project.

## Author contributions

**Conceptualization.** TV, YS, ANJ, ZM, YRS, GM, SS, DL, CL and PKS

**Formal Analysis.** TV, YS, EN, SP, GW, YRS

**Funding acquisition** SS, DL, PKS

**Investigation** TV, YS, SP, RP, ZM

**Supervision** YRS, GM, SS, DL, CL and PKS

**Writing** TV, YS, JBT, PKS

## DATA AND SOFTWARE AVAILABILITY

GeoMx gene expression data will be available via the Gene Expression Omnibus (GEO; RRID:SCR_005012). All CyCIF images with associated histopathological and GeoMx MR annotations will be available to explore online via Minerva software^96^, which supports zoom, pan, and selection actions without requiring the installation of software or downloading the images, at the Harvard Tissue Atlas (https://www.tissue-atlas.org/). **Supplementary Video S1** gives an overview of how to navigate between channel groups and visualize annotations using Minerva. Selected CyCIF channels for specimen MEL14 can be explored via Minerva at https://www.cycif.org/data/vallius-2025/early_melanoma_tumor_intrinsic_e41/MEL14-A3/index.html. Full-resolution CyCIF images, single cell segmentation masks, and cell count tables will be available via the NCI Human Tumor Atlas Network data portal (https://data.humantumoratlas.org/); see **Supplementary Table S1** for a mapping of sample numbers to HTAN IDs. Note that individual image files are ∼100 GB in size. Code used for multimodal spatial analysis are available on GitHub (https://github.com/labsyspharm/2025-Vallius-Shi-Novikov-melanoma-PCAII ).

## METHODS

### Clinical samples

FFPE specimens of primary cutaneous melanoma were retrospectively identified from the archives of the Brigham and Women’s and Massachusetts General Hospitals (n=62 patients, MEL14 to MEL86; see **Supplementary Table S1** for HTAN identifiers and associated clinical metadata). The samples were collected under Institutional Review Board approval (Primary IRB:2019P003828, Secondary IRB: 21-0656) under a waiver of consent. Two sets of specimens were acquired and deeply annotated: sample set A (n=45) was selected for the presence of multiple disease stages within the same sample with an emphasis on stage I melanoma. Sample set B (n=17) was selected for the presence of later stage II disease; multiple stages were often present in these specimens, but this was not a selection criterion.

Fresh FFPE tissue sections were cut from each tumor block. The first section of each block was H&E-stained and used to annotate histopathological ROIs (**Fig. 1C** and **Supplementary Fig. S1B**). The remaining subsequent sections were used for cyclic multiplex immunofluorescence imaging (CyCIF) experiments to characterize markers of melanoma progression and the features of the immune microenvironment within various stages of melanoma. A fresh cut of the samples was performed for deeper profiling with microregion transcriptomic profiling using GeoMx Digital Spatial Profiling (DSP) technology. The clinical, biospecimen, and imaging-level metadata were all collected following the Minimum Information about Highly Multiplexed Tissue Imaging (MITI) standards^97^.

Based on the melanoma diagnostic criteria, the histopathologic annotations included normal skin (N), Precursor (P), Melanoma *in situ* (MIS), Radial Growth Phase melanoma (RGP), and Vertical Growth Phase melanoma (VGP). These ROIs were further classified and subdivided based on the presence of immune infiltrate (brisk TIL, non-brisk TIL, absent). In the case that a single specimen contained more than one histologic region in each category (e.g., precursor regions on both sides of vertical growth phase melanoma), we performed neighborhood analyses separately, as these regions were not physically adjacent.

### Imaging (H&E and tissue-based CyCIF)

H&E-stained FFPE sections from each tissue block were imaged using a CyteFinder slide scanning fluorescence microscope (RareCyte Inc.) with a 20×/0.75 NA objective with no pixel binning. Serial FFPE sections (5 μm thick) were subjected to whole-slide, subcellular resolution, 34 to 49-plex CyCIF imaging with two different antibody panels to generate complementary datasets (**Fig. 1C** and **Supplementary Fig. S1B, S2B**). CyCIF panels 1 and 3 were used for general immune phenotyping to fully profile the tumor microenvironment, and panel 2 was used to better understand the tumor intrinsic features.

Antibody information and associated RRIDs are listed in **Supplementary Table S1**. CyCIF was performed as described in^98^ and more specifically at protocols.io (10.17504/protocols.io.j8nlkoqbdv5r/v1). In brief, the BOND RX Automated IHC Stainer was used to bake FFPE slides at 60°C for 30 minutes, dewax using Bond Dewax solution at 72°C, and perform antigen retrieval using Epitope Retrieval 1 (Leica) solution at 100°C for 20 minutes. Slides underwent multiple cycles of antibody incubation, imaging, and fluorophore inactivation. Antibodies were incubated overnight at 4°C in the dark. Before imaging, a 0.15mm single-sided self-adhesive spacer was applied to the slide, followed by wet-mounting of a glass coverslip using 50% glycerol in 1× PBS. This helps to greatly reduce friction during coverslip removal, ultimately preserving the tissue structure between cycles. Images were acquired using the same microscope and objective as the H&E images.

Slides were soaked in 42°C PBS to facilitate coverslip removal, and then fluorophores were inactivated by incubating slides in a solution of 4.5% H_2_O_2_ and 24 mmol/L NaOH in PBS and placing them under an LED light source for 1 hour. Only antibodies that had passed a multi-step validation process and followed the expected staining pattern were included in downstream analysis.

### CyCIF image pre-processing and quality control

Stitching individual CyCIF images together into a high-dimensional representation for further segmentation and analyses was done using the open-source MCMICRO pipeline^58^ (RRID:SCR_022832), an open-source multiple-choice microscopy pipeline (version:38182748aa0ec021f684ce47248c57340d2f4cc7; full codes available on at https://github.com/labsyspharm/mcmicro). Specific parameters used were optimized after iterative inspection of results, specifically focused on performance of the segmentation module to ensure accurate identification of single cells (params.yml files available at https://github.com/labsyspharm/2025-Vallius-Shi-Novikov-melanoma-PCAII). After generating the segmentation masks, the mean fluorescence intensities of each marker for each cell were computed, resulting in a single-cell data table for each acquired whole-slide CyCIF image. The X/Y coordinates of annotated histologic regions on the whole-slide image were used to extract the single-cell data of cells that lie within the ROI range.

Multiple approaches were also taken to ensure the quality of the single-cell data. At the image level, the cross-cycle image registration and tissue integrity were reviewed; regions that were poorly registered or contained severely deformed tissues and artifacts were identified, and cells inside those regions were excluded. Antibodies that gave low confidence staining patterns by visual evaluation were also excluded from the analyses. The quality of the segmentation was assessed, and the segmentation parameters were iteratively modified to improve the accuracy of the segmentation masks.

### CyCIF single-cell phenotyping

We used a gating-based phenotyping approach to classify cells as described previously.^12^ In short, an open-source visual gating tool (https://github.com/labsyspharm/gater) was used to derive gates for each marker. Similar to batch correction, the identified gates for each marker were then used to rescale the single-cell data between 0 and 1, such that the values above 0.5 identify cells that express the marker and vice versa. The scaled single-cell data was then used for cell-type calling. Tools within the SCIMAP^99^ python package (RRID:SCR_024751) were used to then assign phenotype labels to individual cells based on a hierarchical classification of staining patterns (**Supplementary Fig. S2C**, **Fig. 2A,B**, **Supplementary Table S1,4**). The assigned cell types were verified by overlaying the phenotypic labels onto the images.

### Correlation between protein markers and RNA levels

We performed correlation analysis between expression levels of melanoma gene expression, tumor state pathway scores, and CyCIF quantification of MRs across samples. The pairwise Spearman correlation coefficients were calculated based on the log-transformed RNA level, protein expression level, and pathway scores, as well as CyCIF quantification of each GeoMx ROIs. Pairwise correlation was calculated between those categories. Positive correlation was represented by red circles, with stronger correlations shown by darker and large circles. In contrast, negative correlations were represented by blue circles, with stronger negative correlations shown by darker and larger circles. Non-significant or weak correlations are smaller circles or absent circles near zero.

### Spatial entropy and identification of invasion bands

The local neighborhood Shannon entropy was calculated for each tumor cell according to its nearest neighbors defined via a Delaunay triangulation for the whole-slide images. In order to extend the notion of classical Shannon entropy to spatial entropy, spatial information was incorporated by forming a network graph G = (V, E) using the single cells per tissue specimen. The nodes V of the graph were defined according to the spatial location of single cells and edges E were specified via a Delaunay triangulation. Single-cell neighbors were defined by nearest neighbors in the resulting graph structure and edges were weighted by the Euclidean distance between single cells in tissue space. The graph G was also instantiated as a node attribute graph where each node (single-cell) was assigned the corresponding single cell phenotype (see Methods, CyCIF single-cell phenotyping). Numerous extensions and variations of entropy measures have been proposed in the literature.^100,101^ The spatial entropy metric used here extends the intuition of Wang and Zhao^102^ in classifying landscapes in ecological studies whereby the Shannon entropy is augmented by a prefactor that is directly proportional to the intermixing strength and inversely proportional to the distance between nearest neighboring single cells of different phenotypes. The intermixing strength for each phenotype is defined as the cumulative total of the number of nearest neighbor single cells with differing phenotypes across all local neighborhoods. Given the scale differences between microscopy and ecological data, a scaling factor 𝛼𝛼 is used here for visualization purposes. The form of spatial entropy employed in this work (is to the authors’ knowledge applied for the first time in the single-cell microscopy domain) was chosen for its objective and intuitive nature, as well as prior success in discerning spatial structure between varying ecological landscapes. The mathematical form of the spatial entropy 𝐻𝐻 implemented in the single-cell analysis in this manuscript is detailed below:

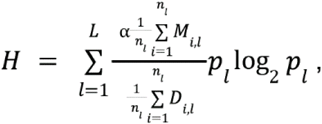

where the intermixing strength 𝑀𝑀_𝑖𝑖,𝑙𝑙_ for cell 𝑖𝑖 with attribute 𝑙𝑙 is defined as,

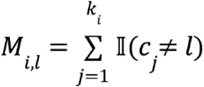

and the distance 𝐷𝐷_𝑖𝑖,𝑙𝑙_ for cell 𝑖𝑖 with attribute 𝑙𝑙 is defined as,

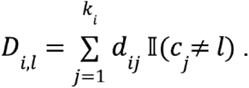

The variables and notation is defined as follows, 𝐿𝐿 is the number of attributes, e.g., single-cell tumor phenotypes, 𝑛𝑛_𝑙𝑙_ is the number of single cells with attribute 𝑙𝑙 in the tissue, 𝑘𝑘_𝑖𝑖_ is the number of nearest neighbors of cell 𝑖𝑖, 𝑐𝑐_𝑗𝑗_ is the attribute of cell 𝑗𝑗, 𝑑𝑑_𝑖𝑖𝑗𝑗_ is the distance between cell 𝑖𝑖 and cell 𝑗𝑗, 𝕀𝕀 is the indicator function which equals 1 if the condition is satisfied and 0 otherwise, 𝛼𝛼 is a scaling factor described above, and 𝑝𝑝_𝑙𝑙_ is the probability of attribute 𝑙𝑙 in the tissue.

An open-source software package SpatialCells^103^ was used to identify subregions with a thickness of 0.2mm based on distance from the epidermis (invasion bands, **Supplementary Fig. S14A**). An increment of 0.2mm was used to separate each subregion. Local progression stages were further split into disjoint spatial subsections (in order not to bias the spatial analysis) and the spatial entropy 𝐻𝐻 was calculated in each subregion.

### Recurrent cellular neighborhood (RCN) analysis

CyCIF neighborhood analyses were performed using the functions *spatial_count* and *spatial_cluster* within the SCIMAP^99^ python package. For each single cell in the CyCIF data, we determined the local neighborhood composition by querying a radius of 50 pixels (15 microns) from the cell centroid as measured by the Euclidean distance between X/Y coordinates. The cell types of these cellular neighbors and their frequencies were identified to generate a neighborhood matrix containing the neighbor phenotype information for every single cell in the data (*spatial_count*). The neighborhood matrix was then clustered using k-means clustering with *k* = 25 (*spatial_cluster*). The optimal number of *K*-means clustering was determined by looking for the elbow point in the computed cluster heterogeneity during convergence (**Supplementary Fig. S13D**). The clusters were then grouped into RCNGs (n=11) based on similarity of the microenvironmental composition. The RCNG assignments were visually confirmed by overlaying the RNCGs on multiple CyCIF images.

### Microregion transcriptomics (GeoMx^Ⓡ^) processing, data collection and annotation

For GeoMx digital spatial profiling (DSP), we used NanoString GeoMx human whole transcriptome atlas (WTA) RNA probes to profile the histopathological annotated regions mentioned above using previously described methods^50^. Briefly, freshly cut (< 2 weeks) 5 μm thick FFPE sections were baked at 60℃ for 3 hours, dewaxed, and hybridized with the WTA probes overnight. On the following day, the slides were incubated with fluorescence-conjugated antibodies targeting melanocytes and tumor cells (SOX10/MLANA), epithelial cells (pan-CK), and immune cells (CD45) before imaging and transcript collection on the DSP (**Supplementary Fig. S1D**, **Supplementary Table S1**). These fluorescent signals were used for ROI selection and for probe collection of 1,266 ROIs representing morphologically distinct sites (N, P, MIS, RGP, VGP; **Fig. 1C**). The collected probes were pooled for library preparation and sent out for Next Generation Sequencing (Biopolymers Facility at Harvard Medical School). In addition to the annotations for local progression axis, the selected GeoMx microregions were annotated for location and immune responses. The annotations for location included epidermis (melanocyte and tumor regions in normal/precursor/MIS and the superficial intra-epidermal component above the RGP and VGP regions), tumor center, stroma, margin (invasive tumor front), and periphery (regions at the periphery of VGP either separate of the main tumor mass, or presenting with discohesive tumor cells at the periphery of VGP without a solid invasive front), tumor center, other). Immune responses for the microregions were annotated by visually assessing the CD45 staining and the cell morphology and were classified as brisk, non-brisk, or absent for RGP and VGP regions, and as immune response or no immune for N, P, and MIS regions.

### GeoMx trajectory analysis

Principal components analysis (PCA) was performed on the tumor (tumor/melanocytic annotated) MRs from all datasets (Sets A1, A2, B1, B2, e.g., **Fig. 3A-I**) without batch correction. Tumor MRs were projected onto the first two principal components (PC), with PC1 and PC2 explaining 28% and 14% of the variance, respectively. The inferred developmental trajectory, depicted as a black line, was computed using the *Slingshot* package (https://github.com/kstreet13/slingshot; RRID:SCR_017012) based on PC1 and PC2 embeddings. In addition, a separate trajectory analysis was performed on Set1 tumor MRs using PCA following batch correction with the *limma* package (e.g., **Fig. 4A**; RRID:SCR_010943).

### Differential gene expression and pathway analysis

Differential gene expression analysis was performed using the *DESeq2* package^104^ (RRID:SCR_015687). The top 60 differentially expressed genes (DEGs) were used for pathway enrichment analysis using the *fgsea* package (https://github.com/alserglab/fgsea) (e.g., in **Fig. 5H**). Additionally, normalized and log_10_-transformed expression matrices extracted from *DESeq2* were used to compute single-sample Gene Set Enrichment Analysis (ssGSEA) scores using the *GSVA* R package (https://www.bioconductor.org/packages/release/bioc/vignettes/GSVA/inst/doc/GSVA.html, RRID:SCR_021058; e.g., **Fig. 8G**).

### Self-organizing map (SOM) and the correlation spanning tree

Gene expression data from GeoMx Set A1 and Set A2 were log10-transformed, quantile normalized and centralized on a gene-wise basis by subtracting the mean log expression of each gene (averaged across all samples) from its observed value. The processed data were then clustered using a self-organizing maps (SOMs) machine learning method^105^, resulting in 900 co-expressed metagenes (SOM was conducted within Set A1 and Set A2 separately due to batch effect issue). The so-called SOM-portrayal method was used to visualize the general gene expression levels of metagenes using a two-dimensional quadratic 30 x 30 pixel map and a maroon-to-blue color code for high-to-low metagene expression values.^105,106^ Local progression stage -specific mean portraits were generated by averaging the metagene landscapes of all cases belonging to one category and difference portraits between them were calculated as the difference between the metagene values in each grid of these maps. Clusters of co-expressed metagenes were identified by selected so-called “cluster areas” in the SOM portraits using overexpression criteria as described previously.^105^ The metagene Clusters (**Fig. 5B** and **Supplementary Fig. S9C,F**) were determined in unsupervised fashion as previously described.^105^

A pairwise correlation analysis (Pearson’s correlation coefficient) was calculated using the metagene expression profiles among all tumor ROIs. The correlation matrix was then used to construct a correlation spanning tree, where each node represents a ROI and the edges represent the correlation strength between the ROIs. Based on the location of the nodes in the correlation spanning tree, each ROI was assigned a Branch ID (Branch 1, 2, or 3).

### Statistical Tests

All statistical tests to infer *P* value for significant differences between groups were performed using the Mann-Whitney U rank test with either *mannwhitneyu* function in the *scipy* Python package (RRID:SCR_008058) or *stats* and *ggpubr* R packages. The difference in the spatial entropy score variance within Breslow depth categories was tested with the Levene’s test using *car* R package.

