## Supplementary Figures for "Spatial determinants of tumor cell dedifferentiation and plasticity in primary cutaneous melanoma"

Supplementary Figure S1. (Related to Fig. 1)

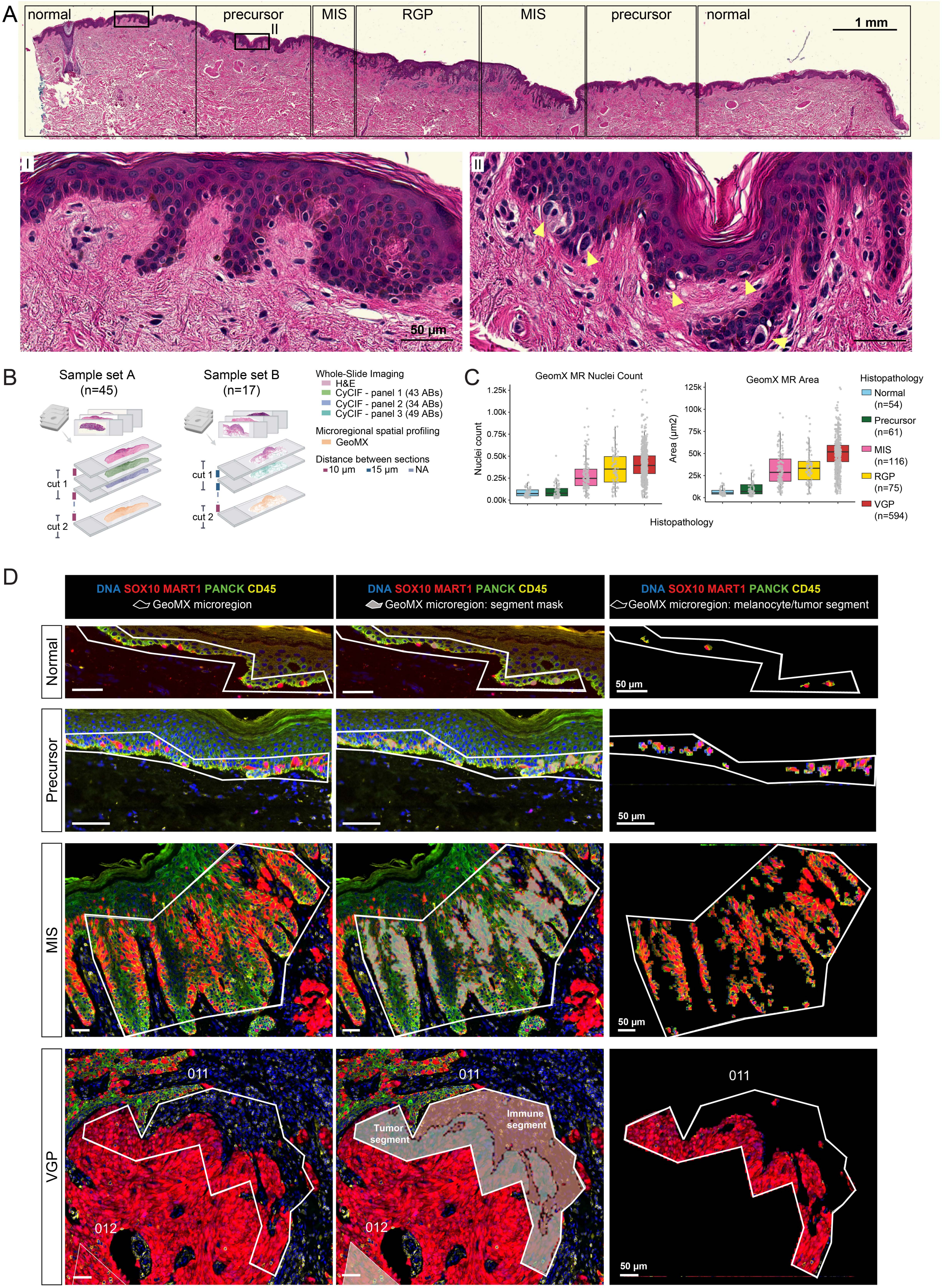

**Supplementary Figure S1 (Related to Fig. 1)**

**A.** Representative example of a primary cutaneous melanoma tissue section stained with H&E (top). The insets I and II represent the magnified regions below, which are examples of melanocytes within normal and precursor regions. Yellow arrows highlight atypical melanocytes within the region of melanocytic atypia. Scale bars, 1 mm and 50 µm. **B.** Schematic figure of sample processing for H&E staining, multiplex tissue imaging (CyCIF) and microregion transcriptomics (GeoMx), illustrating the integration of CyCIF and GeoMx data from adjacent 5 µm sections, guided by histopathological annotations. The GeoMx microregions were mapped to CyCIF images using X/Y coordinates. Each CyCIF panel included 34 to 49 antibodies (see **Supplementary Table S1**). **C.** Box plots showing the number of nuclei (top) and the area (µm$^2$, bottom) of the GeoMx microregions (MRs) within each local progression stage (N, Normal; P, Precursor; MIS, melanoma in situ; RGP, radial growth phase; VGP, vertical growth phase). The datapoints represent individual MRs. Box represents the first and third interquartiles of the data, whiskers extend to show the rest of the distribution except for points that are determined to be outliers. **D.** Representative immunofluorescence images illustrating exemplary GeoMx microregions (MRs) from normal, precursor, MIS and VGP regions (left column). The MR segmentation masks using morphology markers SOX10, MART1 and CD45 are shown in the middle column. The resulting segmented melanocyte/tumor segments are highlighted in the right column. All panels show staining for DNA (blue), SOX10 and MART1 (red), panCK (green), CD45 (yellow). Scale bars, 50 µm.

Supplementary Figure S2. (Related to Fig. 1)

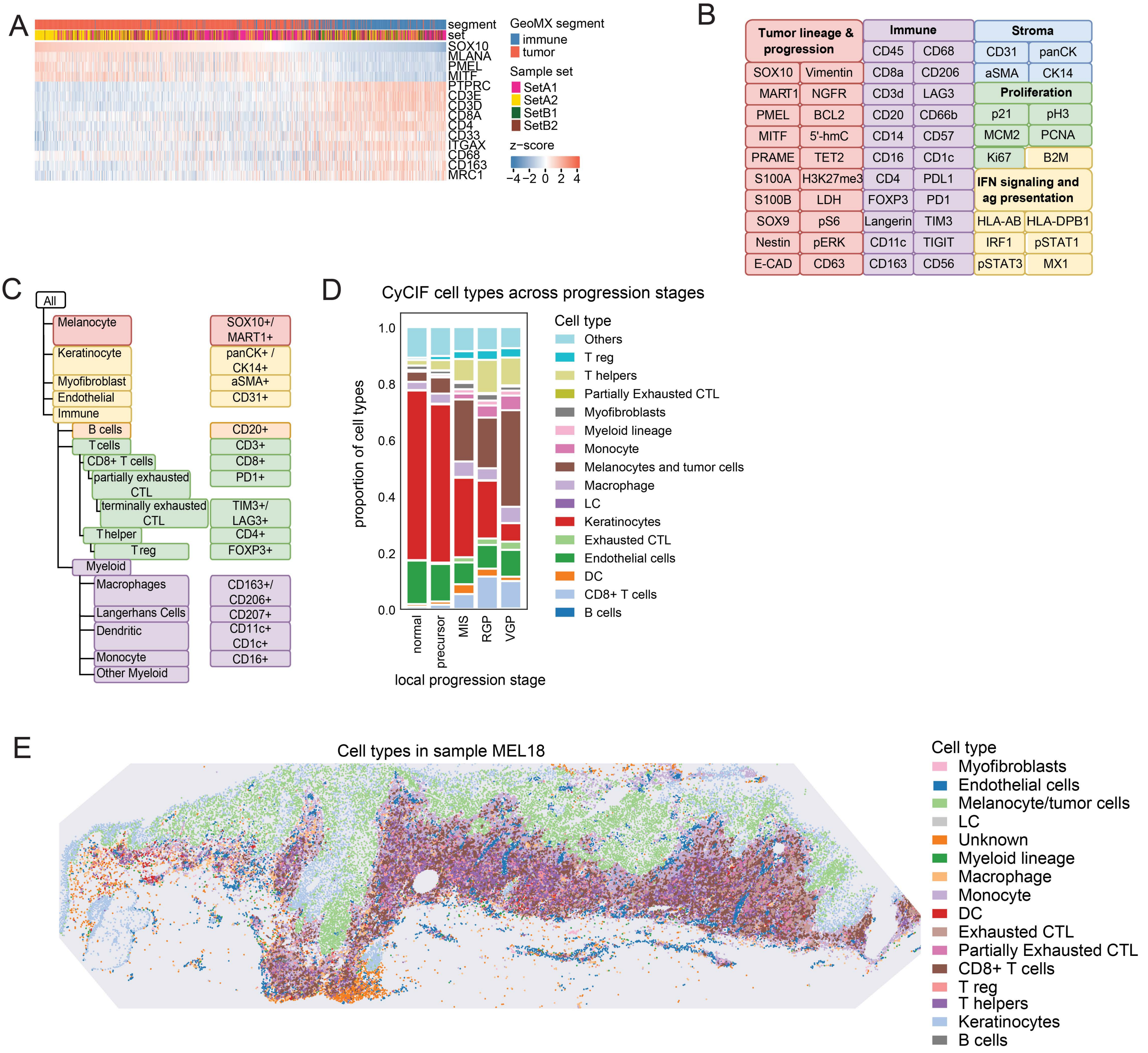

**Supplementary Figure S2 (Related to Fig. 1)**

**A.** Heatmap showing the expression of tumor and immune related genes within GeoMx MR segments. **B.** Antibodies included in CyCIF panels 1-3. All the antibodies used in the experiments are also listed in **Supplementary Table S1** with additional details. **C.** Flowchart of the cell type calling strategy used in CyCIF panel 1 indicating strategy for cell type calling. CyCIF marker intensity data were gated on a per specimen basis and patterns of positive and negative staining of 20 cell lineage and functional markers were used to classify cell types, including four major non-immune cell types (melanocytic, keratinocytic, myofibroblast, and endothelial), B cells, five subtypes or functional states of T cells, and five subtypes of myeloid cells **D.** The proportion of cell types quantified from CyCIF histologic regions across local progression stages for Set A samples. **E.** CyCIF cell types mapped back to their X,Y coordinates within specimen MEL18. The datapoints within the scatter plot represent individual cell centroids and are colored by their cell type annotation.

Supplementary Figure S3. (Related to Fig 2)

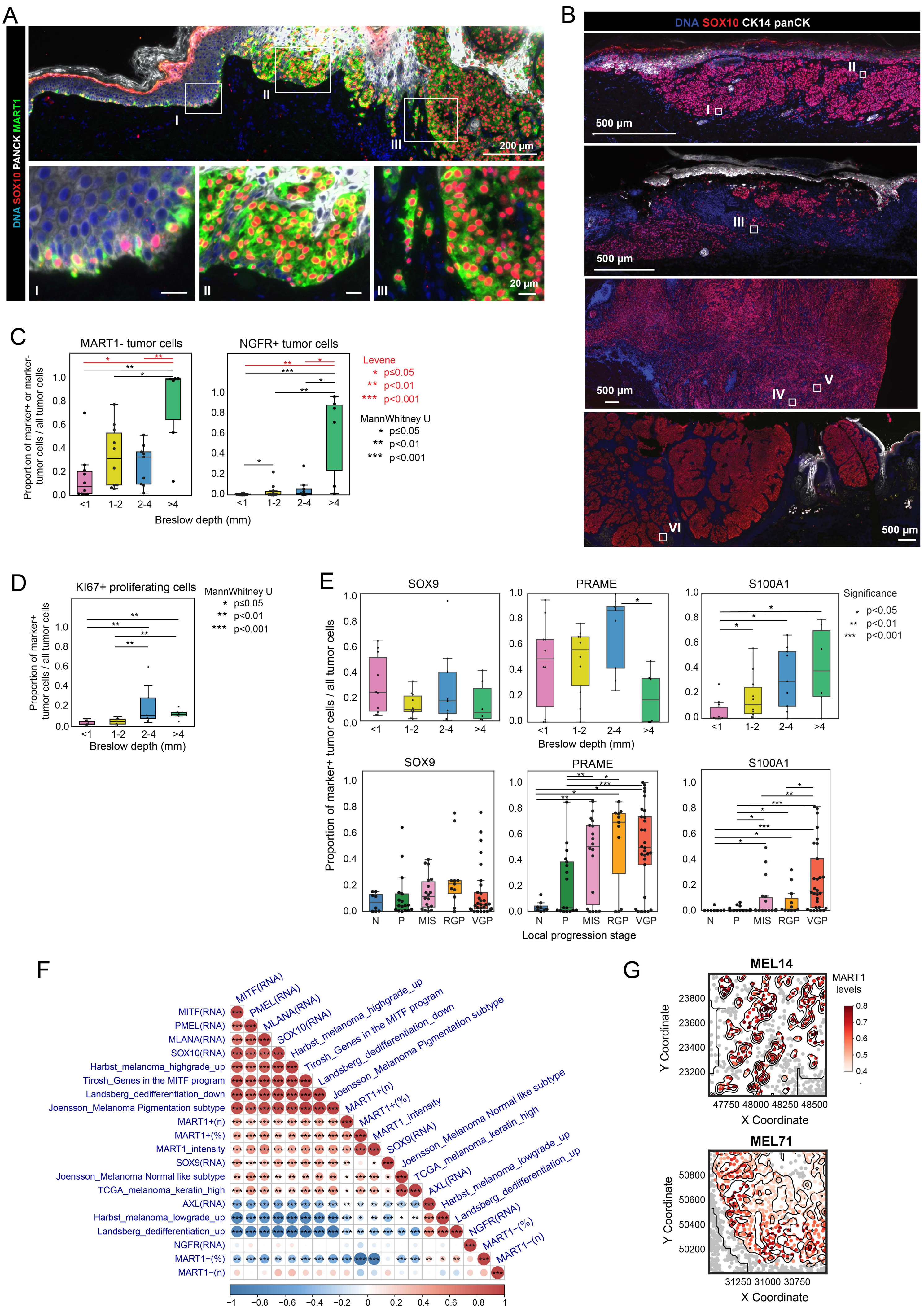

**Supplementary Figure S3 (Related to Fig. 2)**

**A.** Representative fields of view from specimen MEL65. Magnified insets represent regions of precursor (I), MIS (II) and VGP (III). The specimens are stained for DNA (blue), SOX10 (red), MART1 (green) and panCK (white). Scale bars, 200 µm and 20 µm. **B**. Whole-slide images of specimens MEL53, MEL25, MEL82 and MEL71 (from top to bottom), stained for DNA (blue), SOX10 (red), CK14 (white). The ROIs labeled as I-VI are shown in **Fig. 2B**. **C.** The proportion of SOX10+ melanocytes or tumor cells that are MART1-negative (left) and NGFR-positive (right) within tumors with available CyCIF data (whole cohort) grouped by Breslow depth (<1mm, 1-2mm, 2-4mm and >4mm). The datapoints represent mean proportions of marker+ or marker- tumor cells within individual samples. MannWhitney U test (black) and Levene's test (red) *, $p < 0.05$; **, $p < 0.01$; ***, $p < 0.001$. **D.** The proportion of KI67+ tumor cells within tumors with available CyCIF data (whole cohort) grouped by Breslow depth. The datapoints represent mean proportions of marker+ tumor cells within individual samples. MannWhitney U test *, $p < 0.05$; **, $p < 0.01$; ***, $p < 0.001$. **E.** The proportion of SOX10+ melanocytes or tumor cells expressing SOX9, PRAME and S100A1 within tumors with available CyCIF data (whole cohort) grouped by Breslow depth (<1mm, 1-2mm, 2-4mm and >4mm) and local stages of progression (N, P, MIS, RGP and VGP). The datapoints represent mean proportions of marker+ tumor cells within individual samples. MannWhitney U test *, $p < 0.05$; **, $p < 0.01$; ***, $p < 0.001$. **F.** Correlation matrix of gene signatures and phenotype features. Heatmap of pairwise Spearman correlation coefficients among melanoma gene expression, tumor state pathway scores, and CyCIF quantification of MRs across samples. The color of each circle represents the direction and strength of the correlation: red indicates positive correlations, while blue indicates negative correlations, with intensity scaled by the magnitude of the correlation coefficient (range: –1 to 1). The size of each circle reflects the absolute value of the correlation coefficient, while statistical significance is annotated with asterisks: *, $p < 0.05$; **, $p < 0.01$; ***, $p < 0.001$. Significance was assessed using two-sided Spearman correlation tests, and *P*-values were adjusted using the Benjamini-Hochberg false discovery rate (FDR) method. Only significant correlations are annotated. Only the lower triangle of the matrix is shown, and variables are hierarchically clustered to highlight correlated modules. **G**. Scatter plots showing MART1 mean fluorescence intensity for tumor cells within MEL14 (top) and MEL71 (bottom). Fields of view highlighted with rectangles in **Fig. 2H**. The centroids for other cell types are represented with grey dots.

Supplementary Figure S4. (Related to Fig 3)

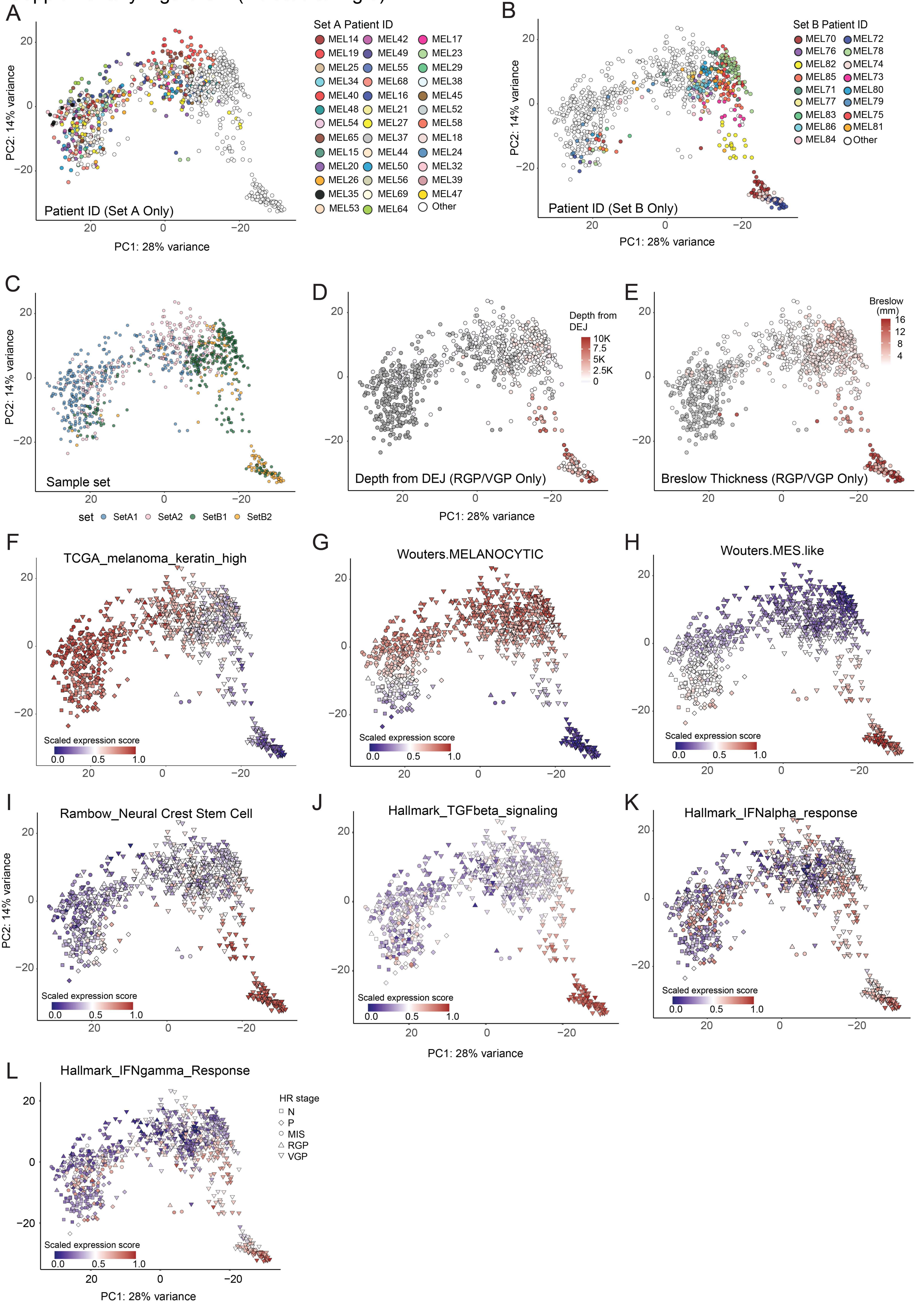

**Supplementary Figure S4 (Related to Fig. 3)**

**A.** Principal component analysis (PCA) plot of GeoMx spatial transcriptomic data from microregions (MRs) representing melanocytes and tumor cells. Each datapoint represents a MR and colors indicate Patient ID for MRs from Set A specimens. **B.** PCA plot of GeoMx spatial transcriptomic data from MRs representing melanocytes and tumor cells. Colors indicate Patient ID for MRs from Set B specimens. **C.** PCA plot of GeoMx spatial transcriptomic data from MRs representing melanocytes and tumor cells. Colors indicate sample set (Set A1, Set A2, Set B1, Set B2). Sample set A (n=45) was selected for the presence of multiple disease stages within the same sample with an emphasis on stage I melanoma. Sample set B (n=17) was selected for the presence of later stage II and III disease; multiple stages were often present in these specimens, but this was not a selection criterion. **D.** PCA plot of GeoMx spatial transcriptomic data from MRs representing melanocytes and tumor cells. Invasive regions are colored by the distance of the MRs from dermo-epidermal junction. Non-invasive (normal, precursor, MIS) regions are colored with grey. **E.** PCA plot of GeoMx spatial transcriptomic data from MRs representing melanocytes and tumor cells. Invasive regions are colored by the diagnostic Breslow depth as annotated for the specimen. Non-invasive (normal, precursor, MIS) regions are colored with grey. **F-L.** PCA plot of GeoMx spatial transcriptomic data from MRs representing melanocytes and tumor cells, colored by the enrichment scores for previously published melanoma-related and Hallmark gene signatures: TCGA_melanoma_keratin_high (**F**), Wouters.MELANOCYTIC (**G**), Wouters.MES.like (**H**), Rambow_Neural Crest Stem Cell (**I**), Hallmark_TGFbeta_signaling (**J**), Hallmark_IFNα response (**K**), Hallmark_IFNɣ response (**L**).

Supplementary Figure S5. (Related to Fig 3)

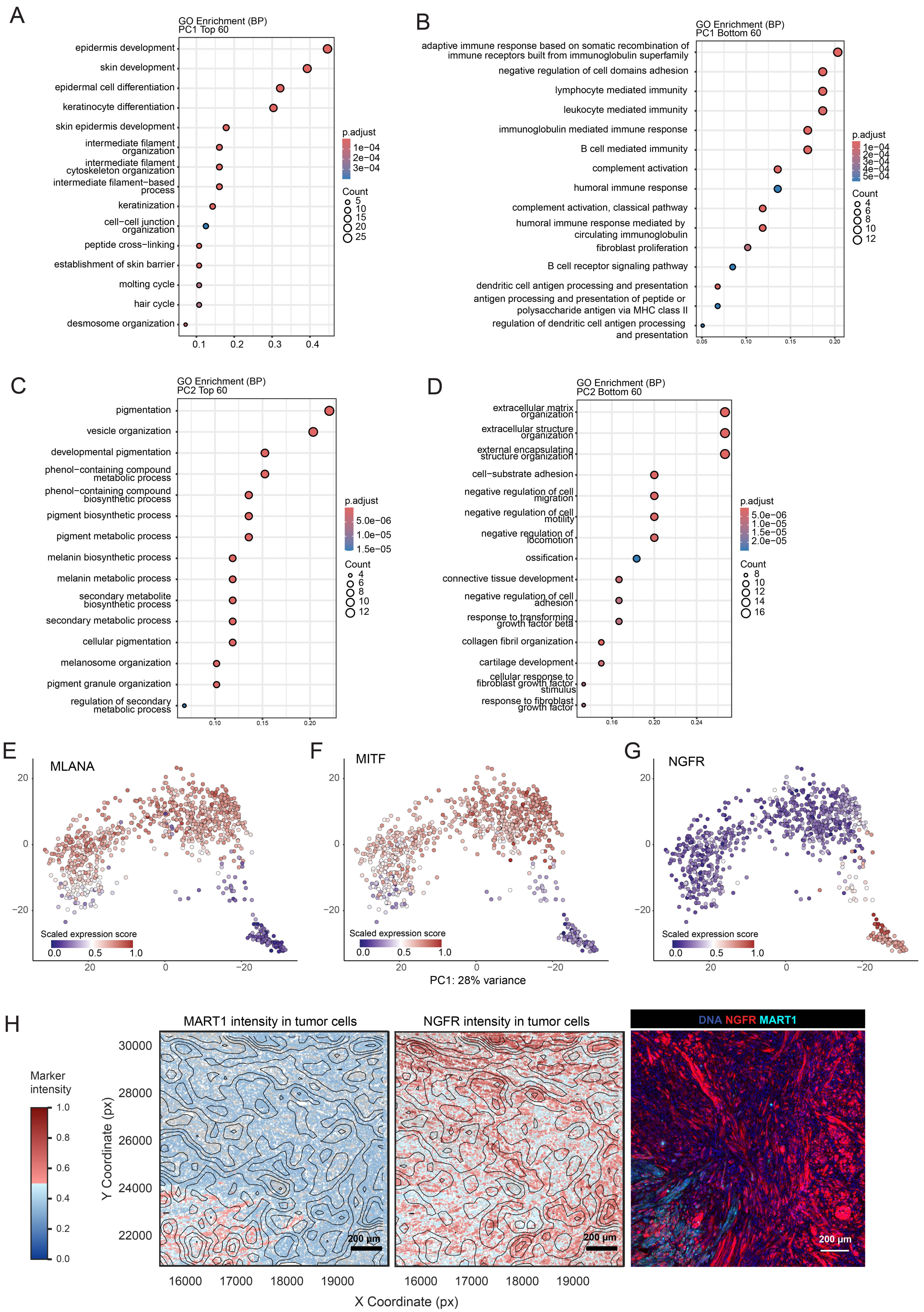

**Supplementary Figure S5 (Related to Fig. 3)**

**A-D.** GO Biological Process (BP) enrichment analysis of the top and bottom 60 genes ranked by PC1 and PC2 loadings. Each dot represents a significantly enriched GO term, with the GeneRatio (number of genes in the gene set overlapping with the GO term divided by the total number of genes in the input list) shown on the x-axis. The size of each dot indicates the number of overlapping genes (Count), while the color represents the adjusted p-value (p.adjust) from multiple testing correction using the Benjamini-Hochberg method. **E-G.** PCA plot of GeoMx spatial transcriptomic data from MRs representing melanocytes and tumor cells, colored by the gene expression levels for *MLANA* (**E**), *MITF* (**F**) and *NGFR* (**G**). **H.** Scatter plots showing MART1 and NGFR mean fluorescence intensities for tumor cells within specimen MEL82. The selected field of view is an expanded view of the magnified region shown in **Fig. 3J**. Color indicates marker intensity (blue to red). The centroids for other cell types are represented with grey dots. Scale bars, 200 μm.

Supplementary Figure S6. (Related to Fig 3)

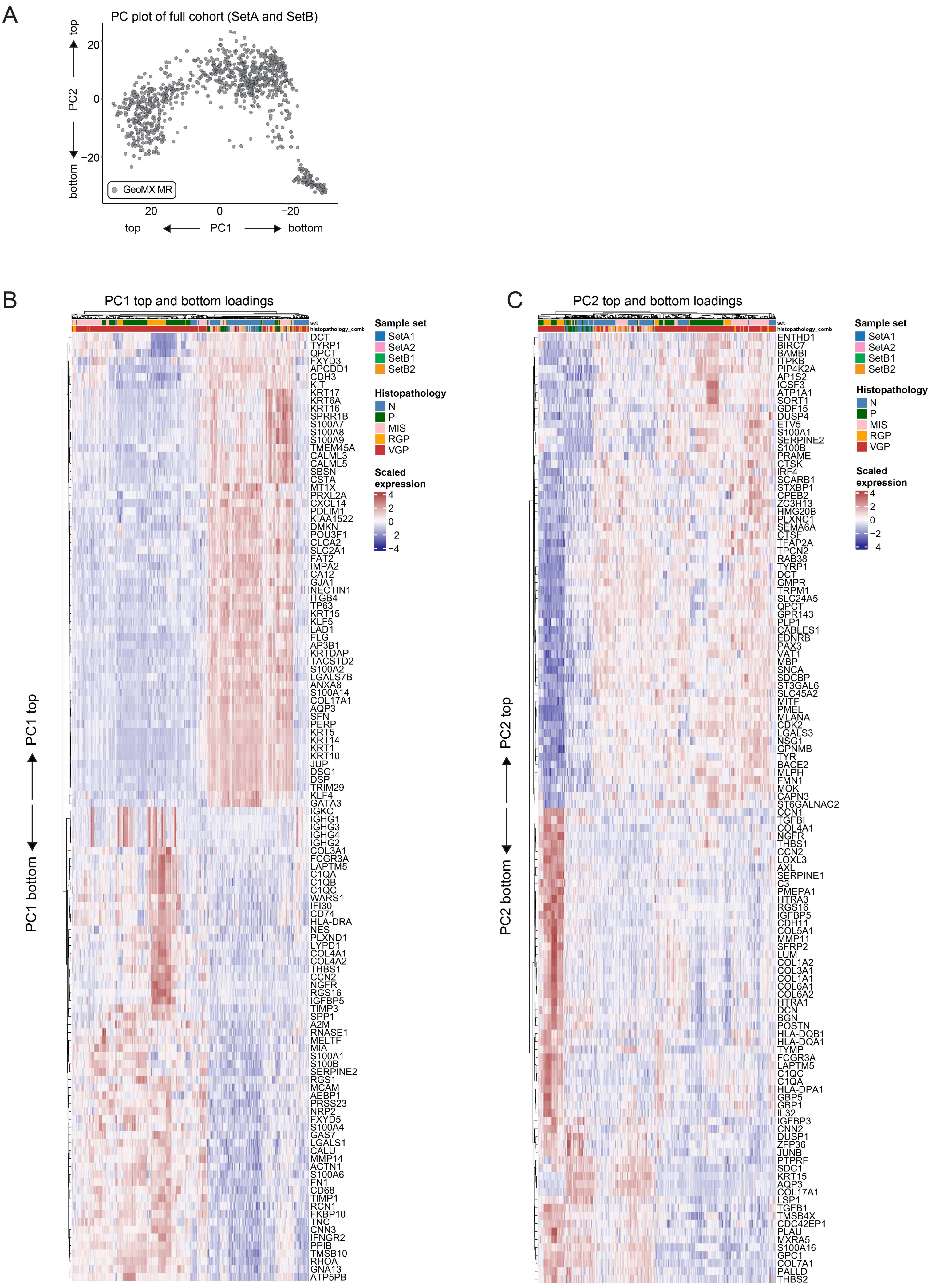

**Supplementary Figure S6 (Related to Fig. 3)**

**A.** Principal component analysis (PCA) plot of GeoMx spatial transcriptomic data from microregions (MRs) representing melanocytes and tumor cells. All MRs are colored with gray. The directions of the axes (top vs bottom) for PC1 and PC2 are shown. **B-C.** Heatmap showing the top and bottom 60 genes contributing to PC1 and PC2. Samples are annotated by Sample set and histopathology. Color scale indicates scaled expression.

Supplementary Figure S7. (Related to Fig. 3)

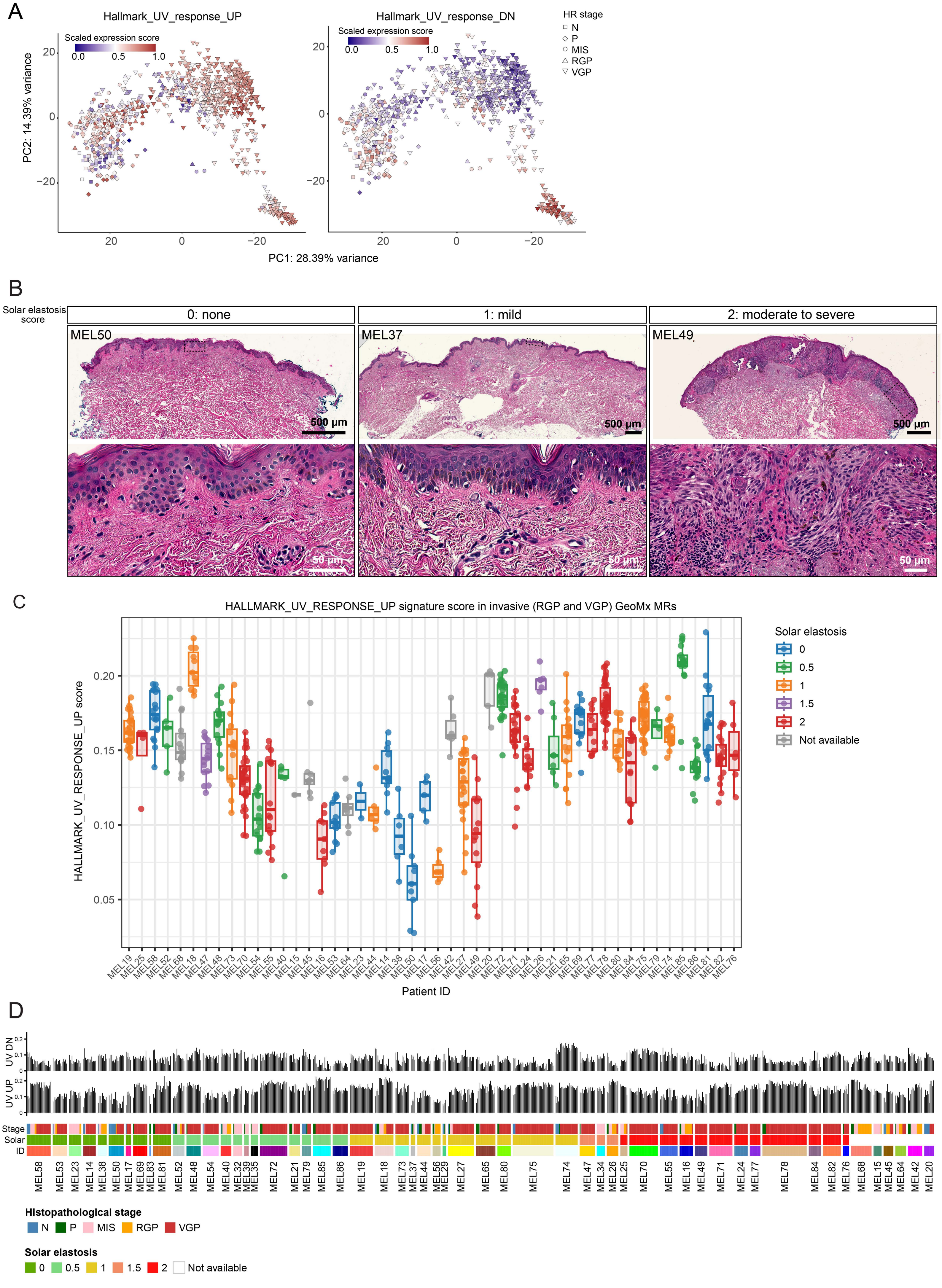

**Supplementary Figure S7 (Related to Fig. 3)**

**A.** Principal component analysis (PCA) plots of GeoMx spatial transcriptomic data from microregions (MRs) representing melanocytes and tumor cells, colored by the enrichment scores for Hallmark_UV_response_UP and Hallmark_UV_response_DOWN signatures. Each datapoint represents an MR and shapes indicate histological region (HR) annotations. **B.** Specimens MEL50, MEL37 and MEL49 annotated for the solar elastosis score (0: none, 1: mild, 2: moderate to severe). **C.** Boxplot showing the Hallmark_UV_response_UP signature scores across patient samples. Each data point represents a gene set expression score calculated for individual invasive (RGP and VGP) GeoMx MRs. Box represents the first and third interquartiles of the data, whiskers extend to show the rest of the distribution except for points that are determined to be outliers. **D.** CoMut plot of the Hallmark_UV_response_UP and Hallmark_UV_response_DN signature scores for each GeoMx microregion (MRs). Each MR is annotated for histopathology (N, P, MIS, RGP, VGP), Solar elastosis score and patient ID.

Supplementary Figure S8. (Related to Fig 4)

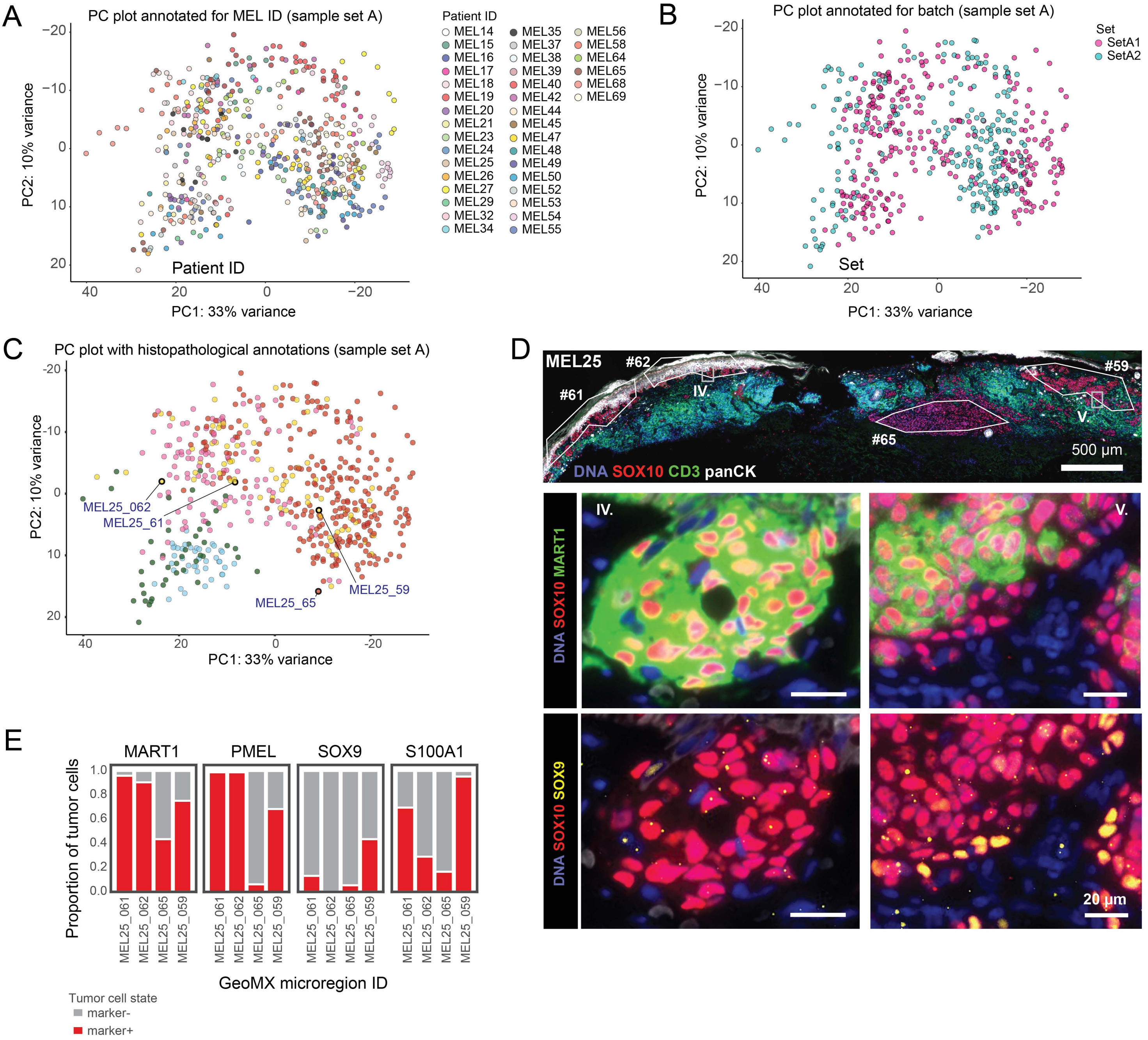

**Supplementary Figure S8 (Related to Fig. 4)**

**A**. Principal component analysis (PCA) plot of GeoMx spatial transcriptomic data from microregions (MRs) representing melanocytes and tumor cells from sample sets A1 and A2. Colors indicate Patient ID for MRs. **B.** PCA plot of GeoMx spatial transcriptomic data from MRs representing melanocytes and tumor cells from sample sets A1 and A2. GeoMx data for sample sets A1 and A2 were collected in separate timepoints, representing two batches of GeoMx data. Colors indicate batch (set A1, set A2). **C.** PC plot of all GeoMx spatial transcriptomic data from microregions (MRs) representing melanocytes and tumor cells for sample Set A, colored by histological annotations. Four exemplary microregions from specimen MEL25 are highlighted (MRs MEL25_061, MEL25_062, MEL25_065, MEL25_059). **D.** Top: CyCIF image of MEL25 labeled with selected VGP GeoMx MRs (white polygons). MRs are labeled with MR ID, indicating how the regions correspond to the PC plot in panel C. Lower: Magnified ROIs from top panel (grey rectangles) showing various tumor cell state markers, including SOX10 (red), MART1 (green), SOX9 (yellow). Scale bars, 500 µm and 20 µm. **E.** The proportion of tumor cells positive for MART1, PMEL, SOX9 and S100A1 within selected VGP GeoMx microregions from specimen MEL25.

Supplementary Figure S9. (Related to Fig 5)

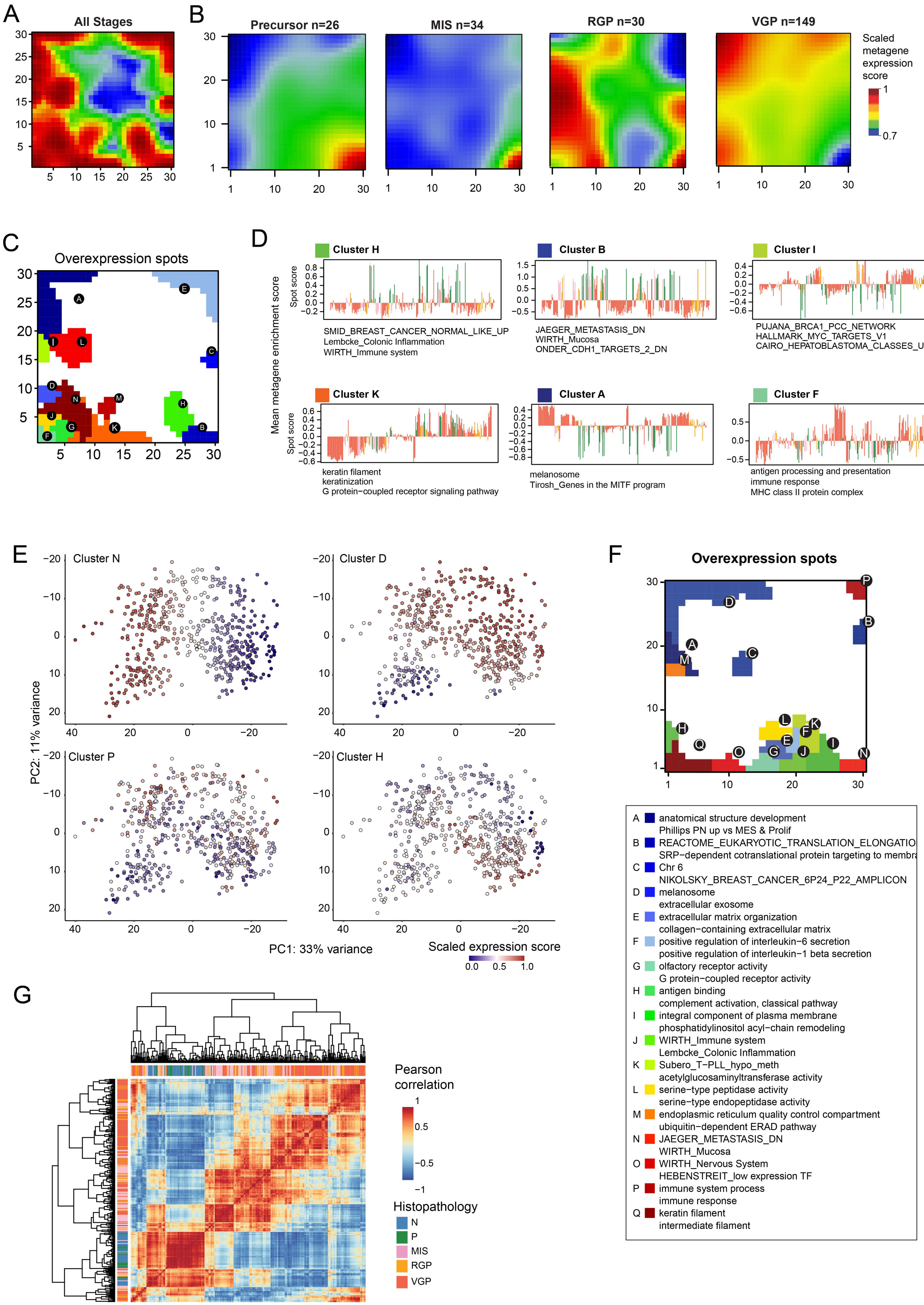

**Supplementary Figure S9 (Related to Fig. 5)**

**A.** SOM portrait of SetA2 GeMX data (n = 239 from n = patients), showing the global expression of 900 metagenes across profiled samples. **B.** SOM portraits for local progression stages (Set A2) showing the global expression of 900 metagenes in MRs grouped by histopathology. The expression levels of each metagene are indicated with a color (scaled metagene expression score) as compared to the average expression levels of all metagenes. **C.** SOM clusters A-N and their mapping to the SOM portrait in **A** (Set A2). The pathways enriched in spots H, B, I, K, A, and F are highlighted in the gray box as they correspond to different histopathological annotations. **D.** Mean expression profiles of the defined SOM clusters H, B, I, K, A, and F within MRs from various progression stage categories (Set A2). The pathways listed below represent the most enriched pathways within each SOM cluster. The Y axis represents the mean expression profile of the SOM cluster signatures of each MR. Individual MRs are shown on the X axis. **E.** PC analysis plot of GeoMx spatial transcriptomic data from Set A from MRs representing melanocytes and tumor cells, colored by the enrichment scores for the overexpression clusters N, D, P and H (defined from Set A1 SOM analysis). **F.** SOM clusters A-Q and their mapping to the SOM portrait for SetA1 analysis. **G.** Heatmap displaying pairwise Pearson correlation coefficients between metagenes for SetA1. The accompanying histopathology annotation indicates the local progression stage most strongly associated with each metagene.

Supplementary Figure S10. (Related to Fig 5)

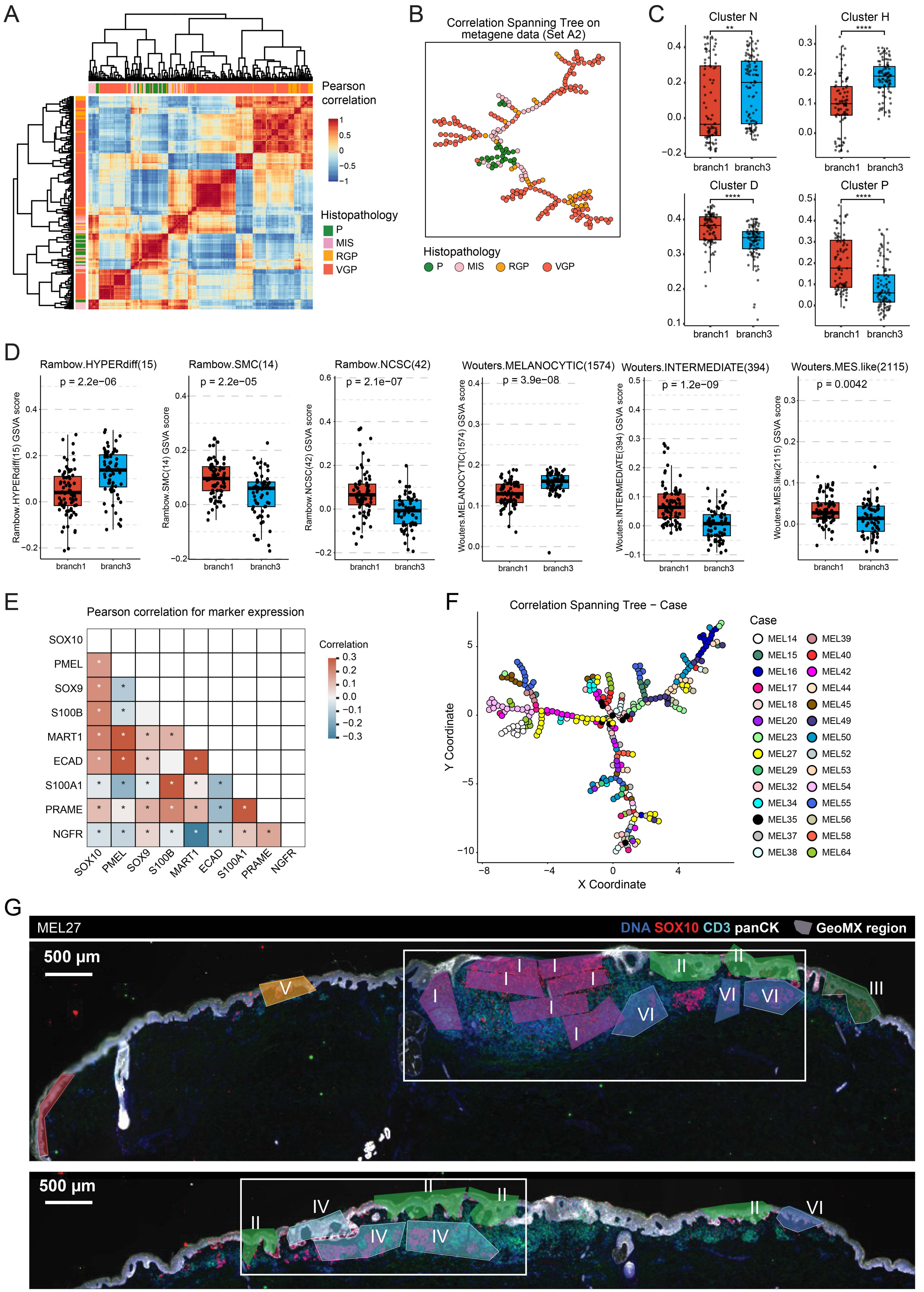

**Supplementary Figure S10 (Related to Fig. 5)**

**A.** Heatmap displaying pairwise Pearson correlation coefficients between metagenes for SetA2. The accompanying histopathology annotation indicates the local progression stage most strongly associated with each metagene. **B.** Minimum spanning tree analysis showing the correlation of profiled GeoMx MRs in SetA2. The minimum spanning tree is annotated by the local progression stage for each MR. Each datapoint represents a single MR. **C.** Boxplot comparison between branch 1 and 3 for overexpression clusters N, H, D and P (Set A1). Each data point represents an expression score calculated for individual GeoMx MRs, and the samples are grouped by the CST branch identity. Box represents the first and third interquartiles of the data, whiskers extend to show the rest of the distribution except for points that are determined to be outliers. The P-value was calculated using the Wilcoxon rank-sum test. **D.** Boxplot comparison between branch 1 and 3 for tumor states (Set A1). Each data point represents a gene set expression score calculated for individual GeoMx MRs, and the samples are grouped by the CST branch identity. Box represents the first and third interquartiles of the data, whiskers extend to show the rest of the distribution except for points that are determined to be outliers. The P-value was calculated using the Wilcoxon rank-sum test. **E.** Matrixplot showing the Pearson correlation of CyCIF marker expression at single-cell level across all the samples in Set A. **F.** Minimum spanning tree analysis showing the correlation of profiled GeoMx MRs in SetA1, colored by Patient ID. **G.** Two sections from specimen MEL27 stained for DNA (blue), SOX10 (red), panCK (white), CD3 (cyan). Colored rectangles indicate MRs and their mapping to the CST in **Fig. 6K**. White rectangles indicate fields of view shown in the CyCIF panels in **Fig. 6K**. Scale bars, 500 μm.

Supplementary Figure S11. (Related to Fig 5)

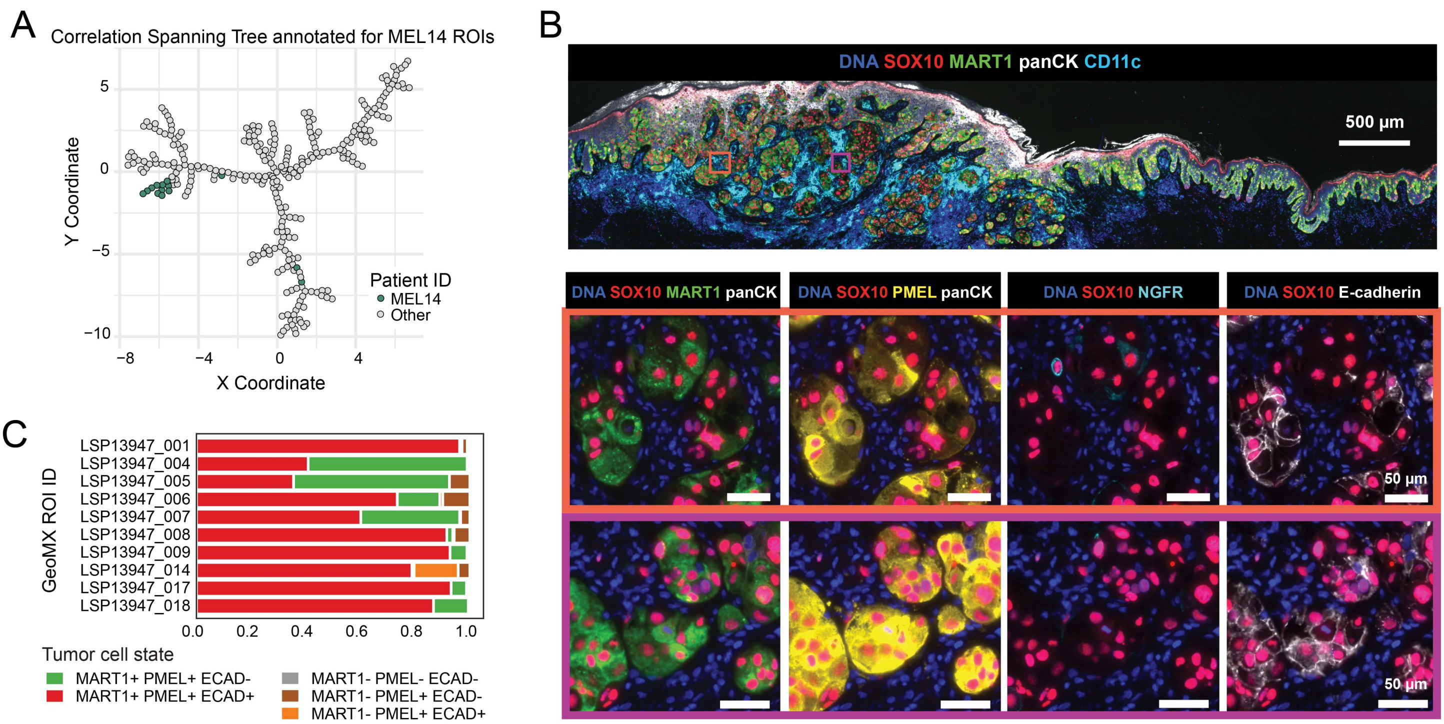

**Supplementary Figure S11 (Related to Fig. 5)**

**A.** Minimum spanning tree analysis showing the correlation of profiled GeoMx MRs in SetA1, colored by MEL14 MRs. **B.** Top: CyCIF image of specimen MEL14 stained for DNA (blue), SOX10 (red), MART1 (green), panCK (white), CD11c (cyan). Bottom: Magnified images of the regions indicated in top, with additional markers: PMEL (yellow), NGFR (cyan), E-Cadherin (white). Scale bars, 500 µm and 50 µm. **C.** Stacked barplot showing the proportion of different tumor states within MEL14 MRs.

Supplementary Figure S12. (Related to Fig 6)

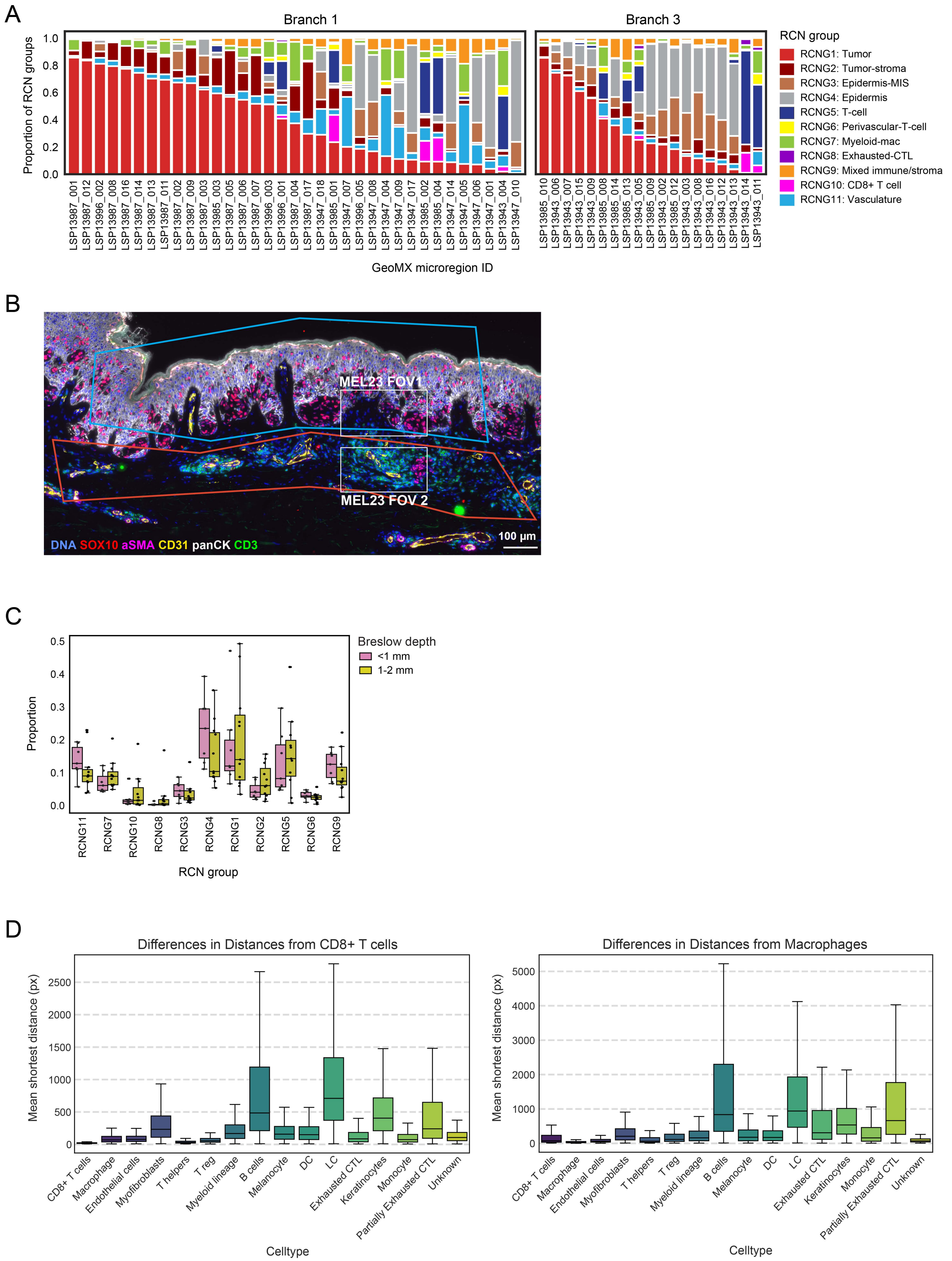

**Supplementary Figure S12 (Related to Fig. 6)**

**A.** The proportion of RCNGs (RCNG1-11) in GeoMx microregions grouped by Branch 1 and Branch 3. **B.** CyCIF image of specimen MEL23, illustrating two MRs belonging to Branch 1 (red rectangle) and Branch 3 (blue rectangle). White rectangles labeled as MEL23 FOV1 and FOV2 are magnified in **Fig. 7G**. **C.** The proportion of RCNGs (RCNG1-11) in whole-slide images grouped by Breslow depth (<1 mm and 1-2 mm). Each datapoint represents the mean percentage of RCNGs within each sample, and the samples are grouped by their Breslow depth (sample-level annotation). **D.** Box plots showing the mean shortest distance from CD8+ T cells (left) and macrophages (right) to all cell types in specimens with CyCIF panel 1 data. Box represents the first and third interquartiles of the data, whiskers extend to show the rest of the distribution except for points that are determined to be outliers.

Supplementary Figure S13. (Related to Fig 6)

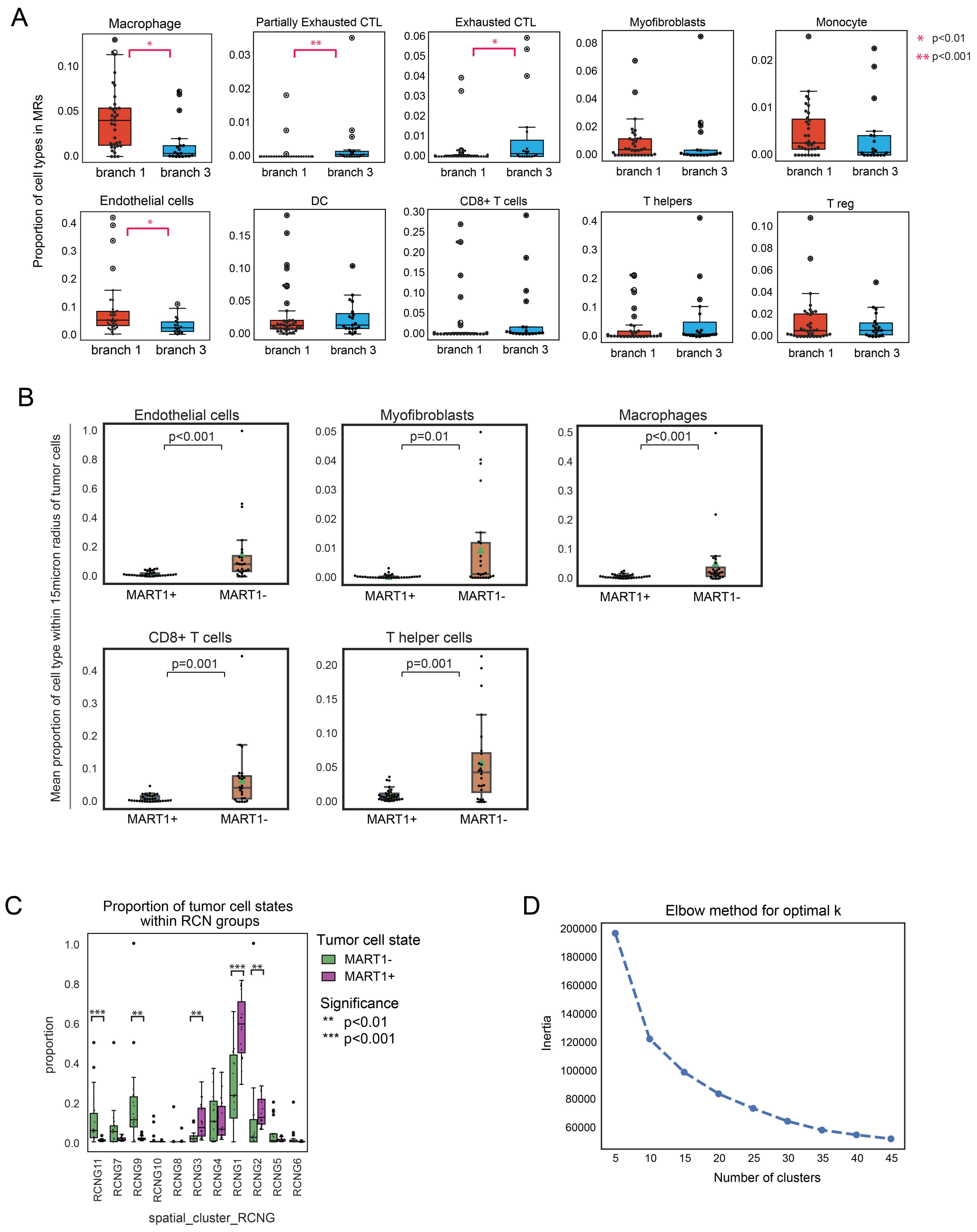

**Supplementary Figure S13 (Related to Fig. 6)**

**A.** The proportion of cell types (partially exhausted CTLs, exhausted CTLs, myofibroblasts, macrophages, monocytes, DCs, CD8+ T cells, helper T cells, T regs) of all cells within Branch 1 and Branch 3 microregions. Each datapoint represents the mean proportion of a cell type within each sample, and the samples are grouped by their correlation spanning tree identity. **B.** The mean proportion of cell types (endothelial cells, aSMA+ cells, macrophages, CD8+ T cells, helper T cells) of all cells within cellular neighborhoods surrounding MART1+ and MART1- tumor cells. Each datapoint represents a mean proportion calculated for a single sample. Box represents the first and third interquartiles of the data, whiskers extend to show the rest of the distribution except for points that are determined to be outliers. **C.** The proportion of MART1+ and MART1- tumor cells belonging to various RCNGs (RCNG1-11) in whole-slide images. Each datapoint represents the mean percentage of MART1+ or MART1- tumor cells belonging to unique RCNGs within a single sample. Box represents the first and third interquartiles of the data, whiskers extend to show the rest of the distribution except for points that are determined to be outliers. MannWhitney U test *, $p < 0.05$; **, $p < 0.01$; ***, $p < 0.001$. **D.** Elbow method for identifying optimal k for K means clustering of the cellular neighborhoods for RCN detection.

Supplementary Figure S14. (Related to Fig 8)

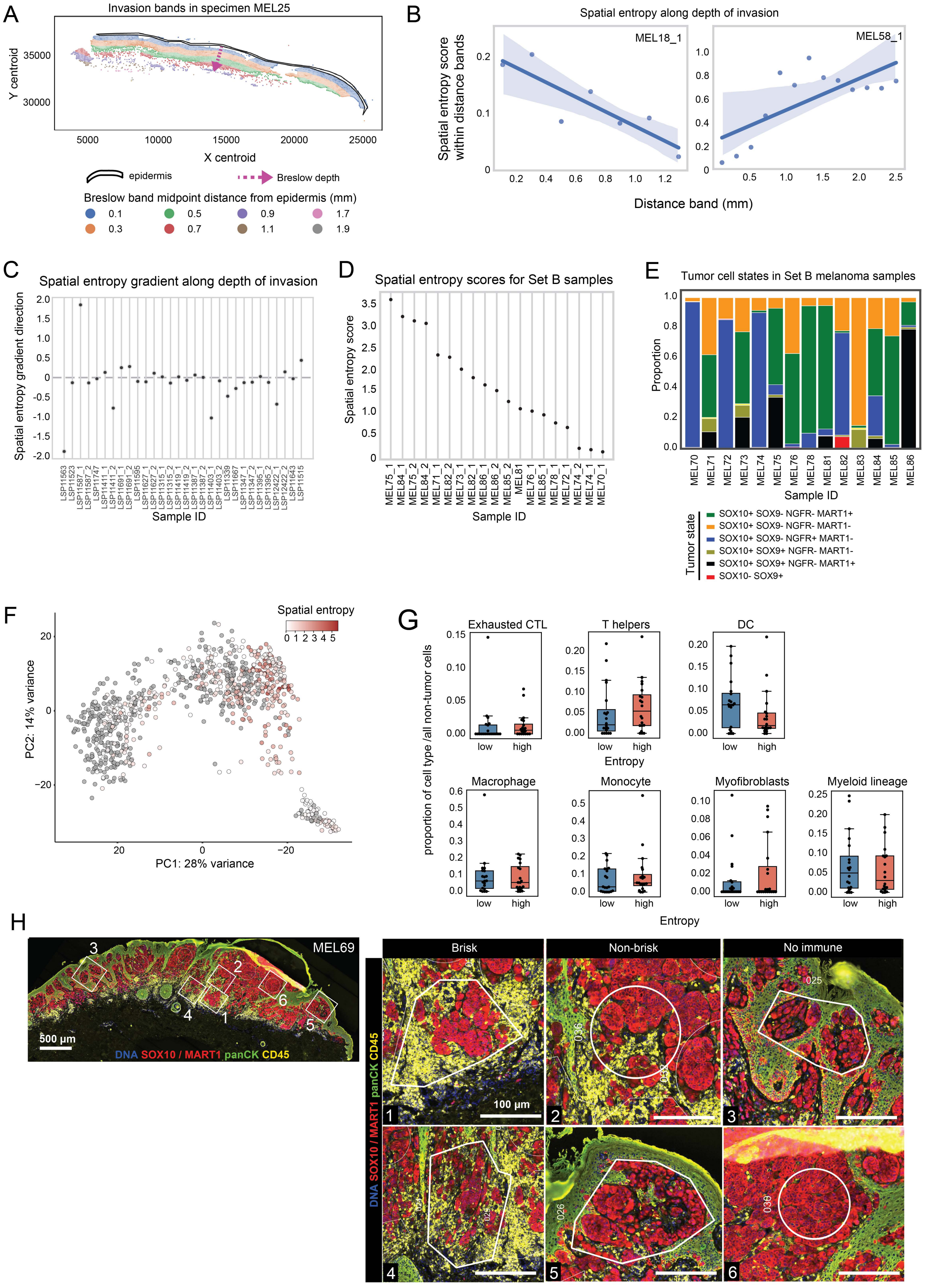

**Supplementary Figure S14 (Related to Fig. 8)**

**A.** Scatter plot of single cell CyCIF data annotated for spatial regions reflecting various depths of invasion ("invasion bands") in specimen MEL25. Each datapoint represents a cell centroid. Purple arrow indicates the measurement of Breslow depth. The spatial regions were separated with an increment of 0.2 mm measured as a distance from epidermis. Colors indicate the midpoint distance of each spatial region. **B.** Correlation plot of spatial entropy score (y-axis) and invasion band (x-axis) of specimens MEL18 (left) and MEL58 (right). **C.** The gradient of spatial entropy score change along the depth of invasion within the profiled tissue sections (Set A samples; see **Methods** for additional details). Note that one specimen can have separate tissue sections that were not spatially connected and hence these regions from the same tissue block were analyzed separately. **D.** Spatial entropy scores for set B samples. **E.** Tumor cell states in set B samples. **F.** PC plot colored by spatial entropy scores as calculated for the GeoMx MRs from Set A and B with available CYCIF data. **G.** The proportion of immune and stromal cells out of all non-tumor cells within high or low entropy microregions (Set A). Box represents the first and third interquartiles of the data, whiskers extend to show the rest of the distribution except for points that are determined to be outliers. **H.** Specimen MEL69 stained for DNA (blue), SOX10 / MART1 (red), panCK (green) and CD45 (yellow). Left: Whole-slide image with ROIs 1-6 indicated and magnified at right. Right: Insets 1-6 below show GeoMx MRs annotated for immune infiltration patterns (brisk, non-brisk, absent) based on CD45 staining. Scale bars 500 μm and 100 μm.
